# EGF and EREG regulate early oncoprotein-mediated tissue transformation in intestinal organoids

**DOI:** 10.64898/2026.11.07.764837

**Authors:** Mario O. Caracci, Birga Soetje, Luis M. Muñoz-Nava, Kirsten Michel, Sabrina Seidler, Petra Janning, Philippe IH. Bastiaens

## Abstract

Somatic mutations, coupled with abnormal extracellular signaling cues, promote tumor development. The abundance and affinity of EGF-family ligands for EGFR influence tissue development and play a central role in organoid culture by differentially regulating the balance between crypt proliferation and villus differentiation in mouse small intestinal organoids (mSIOs).

To test how medium ligand composition affects early oncogenic transformation, we generated a *Ptprg* KO organoid line by CRISPR/Cas9 and two previously described organoid lines harboring synthetic bio-orthogonal expression systems for KRASG12V and β-Ctn4A, activated by the membrane-diffusible ligands TMP or TMP-Cl.

Measurements of crypt and villus region area showed that acute oncogenic activity did not alter organoid growth in media containing 50 ng/mL EGF; however, organoids grown without EGF consistently showed differential growth of the crypt region. Furthermore, EREG and sub-saturating EGF (0.5-5 ng/mL) treatments differentially affected crypt/villus development in each organoid line. Finally, we measured organoid morphology and found that crypt domain spatial distribution is independent of crypt area and requires EGF or EREG, highlighting the role of both proliferation and crypt organization in oncogenic tissue transformation.

**Summary:** (Acute) Oncogenic activity and added EGF or EREG in the culture media modulate tissue transformation in mouse small intestinal organoids by shifting the proliferation/differentiation balance and altering organoid morphology.

## Introduction

Mutations affecting Wnt/β-catenin signaling (Yu et al., 2021) and the EGFR/KRAS/MAPK axis (Drosten and Barbacid, 2020) are found throughout oncogenic malignancies. Particularly in intestinal malignancies, mutations in Adenomatous Polyposis Coli (APC) are observed in nearly 90% of colorectal tumors (Zhang et al., 2016) and are believed to start the adenoma-carcinoma sequence, a process in which a benign polyp acquires additional mutations and becomes an invasive carcinoma (Mizutani et al., 2024). APC is part of the destruction complex regulating the stability of β-catenin, a scaffold protein located in the plasma membrane (PM) associated to adherens junctions and in the nucleus associated with transcription factors in the TCF/LEF family, driving transcriptional activity in response to the presence of Wnt ligands and/or GSK-3β inhibition (Valenta et al., 2012). As a central node in Wnt signaling, β-catenin has been the target of multiple studies in both development and disease (Caracci et al., 2021; Clevers and Nusse, 2012; De Ferrari and Moon, 2006).

Another commonly mutated gene throughout the adenoma-carcinoma sequence is *KRAS* (Walther et al., 2009), a small GTPase located at the PM and acting downstream of receptor tyrosine kinases such as EGFR (Schlessinger and Bar-Sagi, 1994). Addition of EGF-family ligands leads to EGFR phosphorylation and recruitment of GEFs, SOS1 and SOS2, which activate RAS-family proteins by GTP loading (Bos et al., 2007), activating Raf and downstream kinases such as MEK and ERK (Terai and Matsuda, 2005). KRAS-activating mutations commonly affect the GTP-binding site, preventing GTP hydrolysis to GDP and leading to a constitutively active protein (Moore et al., 2020). Similarly, mutations affecting EGFR kinase activity, membrane trafficking, or degradation have also been implicated in colorectal and other malignancies (Napolitano et al., 2024; Normanno et al., 2009), and pharmacological inhibition of EGFR membrane localization and activity is commonly exploited in cancer therapy (Stanland et al., 2025).

The advent of gene editing and organoid technology has revolutionized how genetic diseases are studied, enabling the development of *in vitro* models derived directly from living mouse or human tissue, or from induced pluripotent stem cells. In particular, intestinal organoids are the most studied and have provided unparalleled versatility, recapitulating the epithelial cellular diversity and 3D organization of the intestine (Sato et al., 2009; Sato et al., 2011a; Sato et al., 2011b).

Intestinal organoids derived from human and mouse have been successfully used to model the adenoma-carcinoma sequence using CRISPR-Cas9 gene editing (Matano et al., 2015; Li et al., 2014; Takeda et al., 2019). Additionally, inducible oncogene expression, often by doxycycline-responsive promoters, has been extensively used in mouse models, allowing examination of the role of single driver mutations during oncogenic transformation(Mizutani et al., 2017; Riemer et al., 2017). Constitutive or inducible expression of oncogenic β-catenin variants often drives a clear tissue transformation in cultured organoids (Riemer et al., 2017), highlighting the importance of Wnt/β-catenin signaling in early cancer development. On the other hand, expression of other oncogenic drivers such as KRAS G12V mutation is less clear as doxycycline induction only shows minor effects in ERK activation (Brandt et al., 2019)and constitutive expression of KRAS G12D leads to enlarged crypts (Kotani et al., 2021).

While oncogenic KRAS by itself appears to have minor effects in intestinal tissue, reduction of p53 in the intestinal tissue expressing KRAS G12V enhanced oncogenic transformation in mice (Li et al., 2014), highlighting the requirement of further mutations in the adenoma-carcinoma sequence. However, it is interesting to note that KRAS G12D expression in gastric and pancreatic organoids was sufficient to induce oncogenic transformation, and other oncogenic KRAS variants have major effects in pancreatic (Waters and Der, 2018) and lung cancer (Sunaga et al., 2013) suggesting that oncogenic activity is context-dependent, either through cell-intrinsic mechanisms or extrinsic signaling cues from the surrounding tissue.

Concentration and temporal availability of the Epidermal growth factor (EGF) throughout mSIOs organoid development is determinant for balancing proliferation and differentiation in the crypt and villus domains, respectively (Caracci et al., 2026). Commonly used 50 ng/ml of EGF in the organoid culture media greatly reduced PM localization and reduced protein levels of EGFR, likely due to endocytic trafficking to late endosomes and lysosomal degradation (Boretto et al., 2024; Brüggemann et al., 2021; Caracci et al., 2026). On the other hand, growing organoids without EGF led to smaller organoids, with smaller crypt areas but equivalent villus areas compared to EGF-grown organoids (Caracci et al., 2026). Lack of media EGF increased EGFR at the PM and also increased phosphorylation, suggesting that the endocytic regulation of EGFR regulates the balance of proliferation and differentiation in mSIOs (Caracci et al., 2026). EGF family ligands differentially regulate EGFR endocytic cycle and play a preponderant role in both intestinal homeostasis and tumorigenesis; furthermore, several ligands have been shown to be able to replace EGF in intestinal organoid media (Abud et al., 2021). Since EGFR presence at the PM and overall expression are highly dependent on the availability of EGF or other EGF-family ligands, and this regulation affects organoid proliferation and differentiation dynamics, it is possible that culture media used for *in vitro* growth of organoids could be masking not only the effects of oncoprotein expression but potentially the effects of pharmacological inhibitors (Belair et al., 2020; Klein et al., 2025) Our group has recently developed synthetic bio-orthogonal chemically inducible expression systems, based on fusion proteins with ecDHFR^N18T/A19V^ destabilization domains (DD; *E. coli* dihydrofolate reductase) which target the gene products for degradation unless trimethoprim (TMP) is added to the culture media (Iwamoto et al., 2010). These constructs allow controlled expression of genetically encoded oncoproteins at the protein-stabilization and/or subcellular-localization level, enabling acute, temporally controlled expression and avoiding the potential transforming effects of long-term, constitutive oncoprotein expression, ultimately allowing the study of early oncogenic tissue transformation.

While organoids chronically expressing KRASG12V and β-Ctn4A showed massive tissue transformation resembling cystic organoids, chemically inducible constructs maintained crypt-villus morphology in culture (Munoz-Nava et al., 2026, preprint; Soetje et al., 2026, preprint). Notably, acute oncoprotein stabilization did not alter organoid size compared with controls in ENR media; however, β-Ctn4A-DD increased crypt size and disrupted Paneth cell organization (Soetje et al., 2026, preprint). Surprisingly, while CID-DD-KRASG12V expression did not modify organoid growth and morphology, we found that oncoprotein stabilization increased crypt size in organoids grown in the absence of EGF in the culture media (Munoz-Nava et al., 2026, preprint), suggesting that EGFR PM localization and/or expression levels could be key for early oncogenic KRAS G12V activity To test this idea, in the present work, we examined how acute oncoprotein expression modified organoid development in the presence of saturating and sub-saturating EGF concentrations and the low-affinity ligand Epiregulin (EREG). Additionally, in order to study the effect of oncogenic EGFR activity, we tested the effects of EGF and EREG availability *Ptprg* KO organoids, a membrane-bound tyrosine phosphatase associated to CRC and other cancers (Boni and Sorio, 2022; Duan et al., 2025; Wang et al., 2004) which dynamically removes EGFR phosphorylation in the presence or absence of activating ligands (Joshi et al., 2023; Stanoev et al., 2018). To quantify tissue transformation after 96 h of treatment, we evaluated Crypt and Villus area as markers of proliferation and differentiation, respectively. Additionally, we measured organoid morphology by using a methodology adapted from (Soetje et al., 2026, preprint) based on 10 morphometric features which enable the distinction between normal and aberrant morphologies, as well as distinction of crypt reduction, enlargement or enrichment.

Surprisingly, we found that oncoprotein activity modified crypt development in the absence of EGF or at low concentrations; this observation was consistent across all 3 organoid models. KRASG12V tissue transformation was also apparent in the presence of EREG, a low-affinity ligand that promotes EGFR phosphorylation and endocytic recycling back to the PM (Freed et al., 2017). Finally, *Ptprg* KO organoids grown without EGF showed reduced overall organoid size, with reductions in both crypt and villus domain sizes. Organoid crypt morphology also changed significantly under different culture conditions; notably, these changes were not necessarily correlated with crypt area changes, suggesting that crypt development responds to both intrinsic and extrinsic factors.

Our findings highlight the importance of culture media composition when assessing the potential impact of gene-editing approaches altering oncoprotein expression or the use of acute protein expression systems. While culture media formulations often promote the proliferative state required for tissue buildup and expansion, these signals might mask or dampen the intrinsic activity of cellular components associated with cell fate determination, ultimately impacting the establishment of tissue architecture and morphology.

## Results

### β-Ctn4A-DD acute activity in saturating EGF media leads to enlarged crypt morphology without changing crypt area

mSIOs are normally grown in ENR (EGF, Noggin, R-spondin) media (Sato et al., 2009) and variations of this media composition are widely used to grow organoids derived from different tissues (Mahe et al., 2013; van der Vaart et al., 2021; Wang et al., 2025). We tested organoid development from seeded single crypts in NR media, media without EGF, and media supplemented with either the high-affinity EGFR ligand EGF or the low-affinity ligand EREG. EREG has been previously shown to enhance organoid growth in both mouse and human organoids (Childs et al., 2023; Lemmetyinen et al., 2023), notably EREG treatment enhanced the size of the villus region in mSIOs (Caracci et al., 2026) and was a fundamental driver increasing cellular diversity and inducing crypt-like morphology in human organoids (Childs et al., 2023) making it an interesting ligand to further characterize in organoid development.

To couple EGF-family ligand culture-media composition dynamics with intrinsic, genetically encoded, oncogenic perturbations in mSIOs, we first examined Wnt signaling. In canonical Wnt signaling, in the absence of ligand, endogenous β-catenin is phosphorylated by CK1a and GSK-3β, which are often associated with the destruction complex composed alongside Axin and APC; phosphorylation targets β- catenin for proteasomal degradation (Hart et al., 1998). Wnt ligand addition recruits the kinases and destruction complex to the plasma membrane receptors LRP5/6 and Frizzled, allowing β-catenin to accumulate in the cytoplasm and shuttle to the nucleus, where β-catenin binds to TCF/LEF transcription factors to induce transcription of target genes (Clevers and Nusse, 2012).

Our synthetic bio-orthogonal system (Figure 1A) features a β-catenin fusion protein lacking the phosphorylation sites targeted by CK1 and GSK-3β (β-Ctn4A) (Krishna et al., 2026; Valenta et al., 2012), a fluorescent mCitrine tag (mCit), and a C-terminal destabilization domain (DD, ecDHFR) (Iwamoto et al., 2010), which targets β-catenin for degradation. In the presence of TMP, β-Ctn4A-DD protein is stabilized and accumulates, leading to activation TCF/LEF dependent transcription (Soetje et al., 2026, preprint), ultimately decoupling oncogenic β-catenin activity from Wnt-ligand signaling and endogenous kinase regulation.

**Figure 1.**
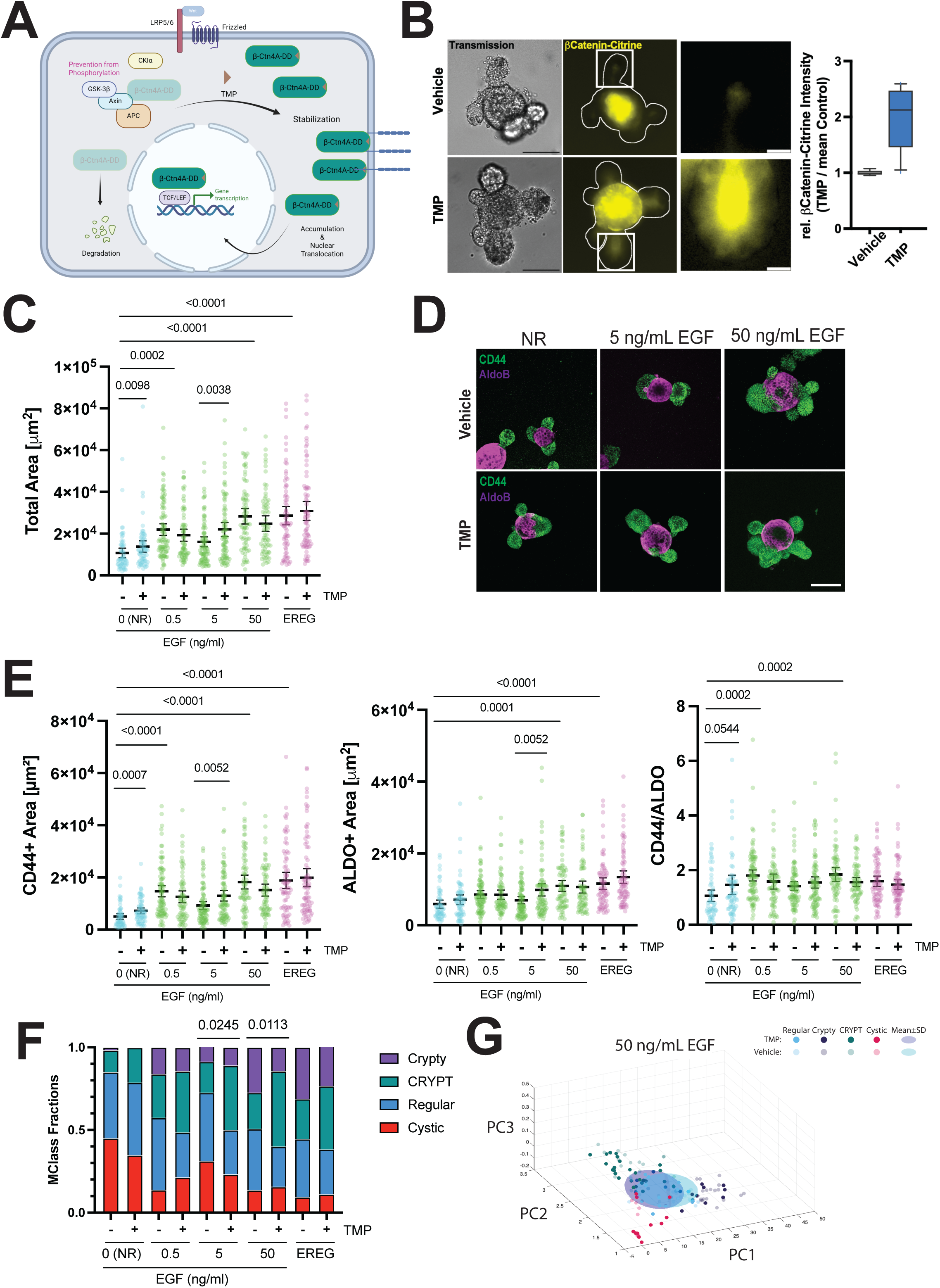
β-Ctn4A-DD acute expression alters crypt/villus growth and crypt domain morphology in NR and lower EGF concentration media. (A) β-catenin4A- mCitrine-DD (β-Ctn4A-DD in figure and text) lacks the phosphorylation sites recognized by GSK-3β and CK1 (4A) and is not targeted by the destruction complex (APC, Axin) for proteasomal degradation. While Wnt ligand binding to LRP5/6 and Frizzled receptors stabilizes endogenous β-catenin, β-Ctn4A-DD contains a destabilization domain (DD) that targets the protein for degradation. Upon addition of the TMP ligand, β-Ctn4A-DD accumulates in the cytoplasm, nucleus, and adherens junctions. Scheme was created with BioRender.com. (B) (Left)Widefield images of organoids seeded from single crypts and treated with 10 μM of TMP or Vehicle. Images show brightfield transmission and mCitrine signal for β-Ctn4A-DD (Yellow). Scale bar: 100 μm. Zoom-in insets denoted by white squares in the center panel. (Right) Box plot showing intensity of mCitrine signal normalized by the mean intensity of the control (Vehicle) condition. Results are from 7 organoids per condition. (C-G) Single crypts were seeded and grown for 96 h with media containing 0 (NR media), 0.5, 5, and 50 ng/mL of EGF or 50 ng/mL of EREG. β-cat4A-mCit-DD was induced by adding 10 μM of TMP in the culture media. (C) Scatter plots showing mean ± 95% CI of Total organoid area in all treatment conditions. Results were obtained from n=62- 81 organoids in N=3 experiments. Significance between Vehicle (-) and TMP-treated (+) organoids was calculated using Mann-Whitney’s test. Group comparisons to NR significance was calculated using ANOVA with Tukey’s test. p-values <0.05 are shown. (D) Representative images of mSIOs showing tissue transformation upon TMP treatment; 50 ng/mL EGF organoids are shown as controls. Organoids were immuno- stained with CD44 (Green), AldoB (Magenta). Scale Bar 100 μm. (E) Scatter plots showing mean ± 95% CI of CD44, ALDOB areas, and the CD44/ALDOB ratio in all treatment conditions. Results obtained from n=62-81 organoids in N=3 experiments. Significance between control (-) and TMP-treated (+) organoids was calculated using Mann-Whitney’s test. Group comparisons to NR significance was calculated using ANOVA with Tukey’s test. p-values <0.05 are shown. (F) Classification of organoid morphologies obtained from n=62-81 organoids in N=3 experiments as described above. Stacked bar plots of morphological Class (MClass) distribution. ‘Regular’ (blue) represents the prototypical crypt-villus structure; versus the aberrant classes: ‘Cystic’ (red) featureless, round organoids; ‘CRYPT’ (green) of enlarged crypts and ‘Crypty’ (violet) with enrichment of crypts. P-values: Chi^2^-test, p-values <0.05 are shown. (G) Representation of organoid morphologies in the morphometric space (principal components PC1, PC2, PC3). Representative PC of significantly altered morphologies, more examples in Figure S4. TMP-treated organoids and the mean±SD ellipsoid (solid colors; purple) versus vehicle-treated and the mean±SD ellipsoid (pastel colors; aqua), visualizing the shift of morphological class distribution towards enlarged crypts (‘CRYPT’, green) in organoids were β-Ctn4A-DD was stabilized by TMP (10 µm) in 50 ng/mL EGF media condition.

Treatment of β-Ctn4A-DD-expressing organoids for 96 h with 10 μM TMP led to a 2- fold increase in total fluorescent signal; the higher intensity is often located in the central villus region where the differentiated, non-proliferative cells are located, with lower-intensity protein stabilization observed in the proliferating crypt area (Figure 1B)

To contrast constitutive expression of oncogenic β-Ctn4A with the acute, inducible expression of βCtn4A-DD, we first grew organoids expressing βCtn4A with 50 ng/mL of EGF, which is usually used in mSIOs culture, and compared it to lower, sub- saturating concentrations of EGF (5 and 0.5 ng/mL and 50 ng/mL EREG. Chronic expression of βCtn4A led to massive modification of organoid morphology, increasing the number of cystic organoids (Soetje et al., 2026, preprint) with widespread expression of CD44 and almost no expression of differentiation markers such as AldoB (Figure S1A and B). Notably, NR, EGF, or EREG treatments did not alter the total area of cultured organoids or enhance AldoB differentiation (Figure S1A and B), suggesting that organoids maintain a persistent non-differentiated precursor state largely independent of environmental growth factors. In sharp contrast, acute, TMP-induced stabilization of β-Ctn4A-DD maintained a differentiated crypt-villus morphology in all treatments (Figure 1D).

Sub-saturating concentrations of EGF are often sufficient to induce organoid growth significantly compared to organoids grown with NR media; this effect is most notably represented in the crypt proliferation, with less effect on villus differentiation (Caracci et al., 2026). Adding TMP did not affect the total area of organoids grown with either 50 ng/ml EGF or EREG; however, it did lead to larger NR-grown organoids (Figure 1C and D). In absence of media EGF, crypt development increased significantly in combination with TMP-induced oncogenic β-catenin activity compared to untreated controls. Furthermore, the CD44/ALDO ratio increased significantly, suggesting that organoid development is altered promoting crypt growth (Figure 1E). While TMP treatment in NR media had no effect on villus development, 5 ng/ml EGF media induced significant growth in both CD44+ and ALDO+ areas while maintaining the CD44/ALDO ratio, suggesting balanced development of both regions (Figure 1E and D).

As observed in our results, organoid development, while stereotypical (Crypt-Villus or Cystic morphology), is highly heterogeneous, with individual organoids showing distinct crypt-region organization, either featuring one or multiple spatially separated crypts or enlarged crypts relative to the organoid size. In principle, this spatial and morphological organization of the crypt domain in developing organoids is determined by proliferation-mediated crypt inflation followed by crypt fission events (Tallapragada et al., 2021), furthermore, crypt dysplasia is a common feature in intestinal tumorigenesis (Olpe et al., 2021; Wasan et al., 1998; Wong et al., 2002).

Finally, our group had previously determined that β-Ctn4A-DD activity modified organoid crypt morphology in the presence of normal organoid culture media with 50 ng/mL of EGF, where TMP addition shifted the percentage of organoids showing a regular morphology, featuring a crypt/villus morphology with distinctly separated crypts, to a morphology characterized by elongated organoids with fewer, enlarged crypts (Soetje et al., 2026, preprint).

In order to asses organoid morphology and crypt organization in an unbiassed way we facilitated a morphometric analysis established in (Soetje et al., 2026, preprint), based on 10 scale- and rotationally invariant morphometric features. These features are shown in the principal component space (PC1, PC2, PC3). While the prototypical crypt-villus morphology as the ‘Regular’ class was determined by the amount of the majority (mean +/- 0.7SD) of vehicle-treated β-Ctn4A-DD organoids grown in ENR, a k-means clustering of all organoids outside this region allowed to determine the aberrant clusters which were fused based on main morphological features of crypt enrichment (PC1) and crypt enlargement (PC2) (See *Methods*).

Morphological classification showed that, regardless of TMP addition, starting at 0.5 ng/mL of EGF or EREG treatments increased the percentage of organoids featuring multiple crypts (“Crypty”, Violet) or enlarged crypts (“CRYPT”, Green) and the reduction of cystic organoids (“Cystic”, Red) compared to organoids grown in NR condition (Figure 1F and G, Figure S4A). Notably, we observe an increased percentage of organoids showing enlarged crypts after TMP treatment at EGF concentrations of 5 and 50 ng/mL (Figure 1F and G), suggesting that crypt enlargements associated with the β-Ctn4A-DD oncoprotein require saturating amounts of EGF, with no significant morphological effects observed in EREG treatment.

While our previous analysis quantified the differential growth of crypt and villus regions of developing organoids, our results highlight that EGF and EREG presence in the culture media are required for the development of crypts in mSIOs, where β-Ctn4A- DD activity enhances a morphological change in crypt organization without necessarily affecting the size (or proliferation) of crypts.

### CID-DD-KRASG12V acute signaling activity increases crypt area but does not modify organoid morphology

For acute chemical activation of KRAS oncoprotein activity, KRASG12V, a KRAS variant locked in an active GTP-bound state (Scheffzek et al., 1997), was split into a membrane-bound hypervariable (HVR) region (TH-tKRAS) and its G-domain lacking the HVR region (CD-DD-GDom), which is located in the cytoplasm. CD-DD-GDom contains the constitutively active G-domain of KRAS, fused to a tandem of two *Escherichia coli* dihydrofolate reductase destabilization domains (DD, ecDHFR) and the mCitrine fluorescent protein (mCit); on the other hand, TH-tKRAS contains the HVR domain of KRAS, fused with a tandem of two modified haloalkane dehalogenase proteins (2XhaloTag) and mTFP fluorescent protein (Figure 2A). Adding TMP-Cl stabilizes and recruits the CD-DD-GDom to the PM, where it activates downstream signaling by Raf and ERK (Muñoz-Nava et al., 2026, preprint). In mSIOs, our synthetic 2-protein signaling system activated by chemically induced dimerization (CID), hereafter referred to as “CID-DD-KRASG12V,” shows constitutive expression of TH- tKRAS associated with the PM and inducible expression of the CD-GDom, which is stabilized and enriched at the PM after 96 h of TMP-Cl addition (Figure 2B). CID-DD- KRASG12V mSIOs showed no transformation phenotype with either DMSO or TMP- Cl treatment (Figure 2D). Contrastingly, Organoids stably expressing mKate2- KRASG12V showed a fully transformed phenotype featuring cystic CD44+ organoids with no expression of AldoB, total area was not affected by NR, EGF or EREG treatment (Figure S1C and D), similar to the observations on chronic expression of βCtn4A.

**Figure 2.**
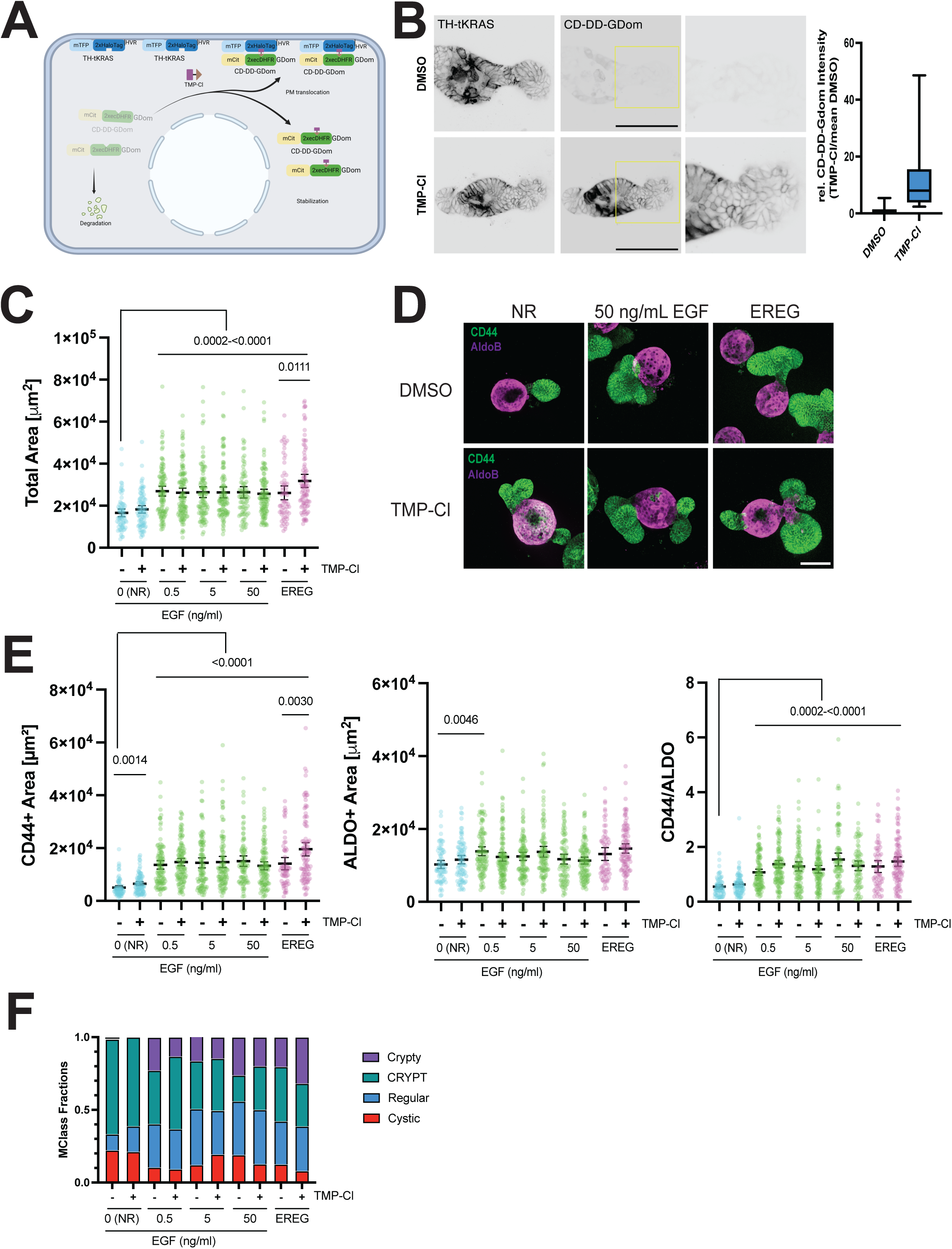
Signaling activity of CID-DD-KRASG12V enhances crypt growth in absence of EGF and in the presence of EREG. (A) Scheme showing from left to right: TH-tKRAS (mTFP-2xHaloTag-HVR) located at the PM and CD-DD-GDom (mCitrine-2xec[DD]DHFR-GDom) in the cytoplasm and destabilized for degradation; upon TMP-Cl addition (TMP [purple rectangle] fused to Cl [brown triangle]; 1 µM), CD- DD-GDom is stabilized and translocated to the PM. The scheme was created with BioRender.com. (B). (Left) Inverted images of TH-tKRAS and CD-GDom of organoids seeded from single crypts and treated with 10 μM TMP-Cl or DMSO. Scale bar: 100 μm. Zoom-in insets are denoted by yellow squares in the center panel. (Right) Box plot showing intensity of CD-DD-GDom signal normalized by the mean intensity of the DMSO condition. Results obtained from 23-29 organoids per condition (C-F) Single crypts were seeded and grown for 96h with media containing 0 (NR media), 0.5, 5, and 50 ng/mL of EGF or 50 ng/mL of EREG. We induced stabilization and membrane recruitment of CD-DD-GDom by adding 10 μM TMP-Cl to the culture media. (C) Scatter plots showing mean ± 95% CI of Total organoid area in all treatment conditions. Results obtained from n= 65-105 organoids in N=3 experiments. Significance between DMSO (-) and TMP-Cl treated (+) organoids was calculated using Mann-Whitney’s test. Group comparisons to NR significance was calculated using ANOVA with Tukey’s test. p-values <0.05 are shown. (D) Representative images of mSIOs showing tissue transformation upon TMP treatment; 50ng/mL of EGF organoids are shown as controls. Organoids were immuno-stained with CD44 (Green), and AldoB (Magenta). Scale bar: 100 μm. (E) Scatter plots showing mean ± 95% CI of CD44, ALDOB areas, and the CD44/ALDOB ratio in all treatment conditions. Results obtained from n=65-105 organoids in N=3 experiments. Significance between DMSO (-) and TMP-Cl-treated (+) organoids was calculated using Mann-Whitney’s test. Group comparisons to NR significance was calculated using ANOVA with Tukey’s test. p-values <0.05 are shown. (F) Classification of organoid morphologies obtained from n=65-105 organoids in N=3 experiments as described above. Stacked bar plots of morphological Class (MClass) distribution. ‘Regular’ (blue) represents the prototypical crypt-villus structure; versus the aberrant classes: ‘Cystic’ (red) featureless, round organoids; ‘CRYPT’ (green) of enlarged crypts and ‘Crypty’ (violet) with enrichment of crypts.

We had previously observed that CID-DD-KRASG12V organoids showed no changes in growth or morphology in the presence of EGF; however, a significant increase in crypt size could be observed when organoids were grown in NR medium (Muñoz-Nava et al., 2026, preprint). Consistent with this, we observed a significant increase in CD44+ area in TMP-Cl-treated organoids grown without EGF; in the presence of EGF, we saw no differences at any tested concentrations (Figure 2C-E). Surprisingly, we observed a sharp, significant change in total and CD44+ area in TMP-Cl-treated organoids grown with EREG, accompanied by a slight increase in ALDO+ area (Figure 2C-E), suggesting that oncogenic KRASG12V activity is mostly associated with crypt domain proliferation.

Morphological classification of CID-DD-KRASG12V showed that adding EGF or EREG to the culture media reduced the percentage of cystic organoids and increased the percentage of multiple-crypt organoids, regardless of TMP-Cl presence (Figure 2F, Figure S4B). While NR and EREG media significantly increased crypt area, we were unable to detect significant changes in organoid crypt morphology in response to acute oncoprotein activity (Figure 2F, Figure S4B), suggesting that, under specific culture media conditions, CID-DD-KRASG12V enhances crypt proliferation but does not selectively modify crypt morphology.

### EREG alters organoid morphology in *Ptprg* KO organoids while NR and sub- saturating EGF alters crypt/villus growth

EGFR phosphorylation is tightly regulated by a spatio-temporally defined network of phosphatases, where the plasma membrane-bound protein tyrosine phosphatase receptor gamma (PTPRG) and ER-bound PTPN2 maintain cells receptive to evolving concentrations of EGF in the cellular environment (Joshi et al., 2023; Stanoev et al., 2018). Gene editing of *ptprg* in MCF7 shows increased phosphorylation of EGFR and downstream targets such as AKT and ERK; notably, these cells also showed constitutive pro-migratory behavior in the absence of EGF in media compared to WT cells(Joshi et al., 2023). *Ptprg* (NM_008981) is a 30-exon gene with 3 documented transcript isoforms (Taki et al., 2024). Its phosphatase activity is dependent on a cytosolic D1 domain common to the PTP phosphatase family, which contains conserved WPD-Loop residues and a catalytic P-Loop Cysteine (Tautz et al., 2013). In order to generate a loss-of-function (LoF) knockout (KO) of PTPRG, we decided to target the complete exon 21, which contains all these critical aminoacidic residues (Figure 3A).

**Figure 3.**
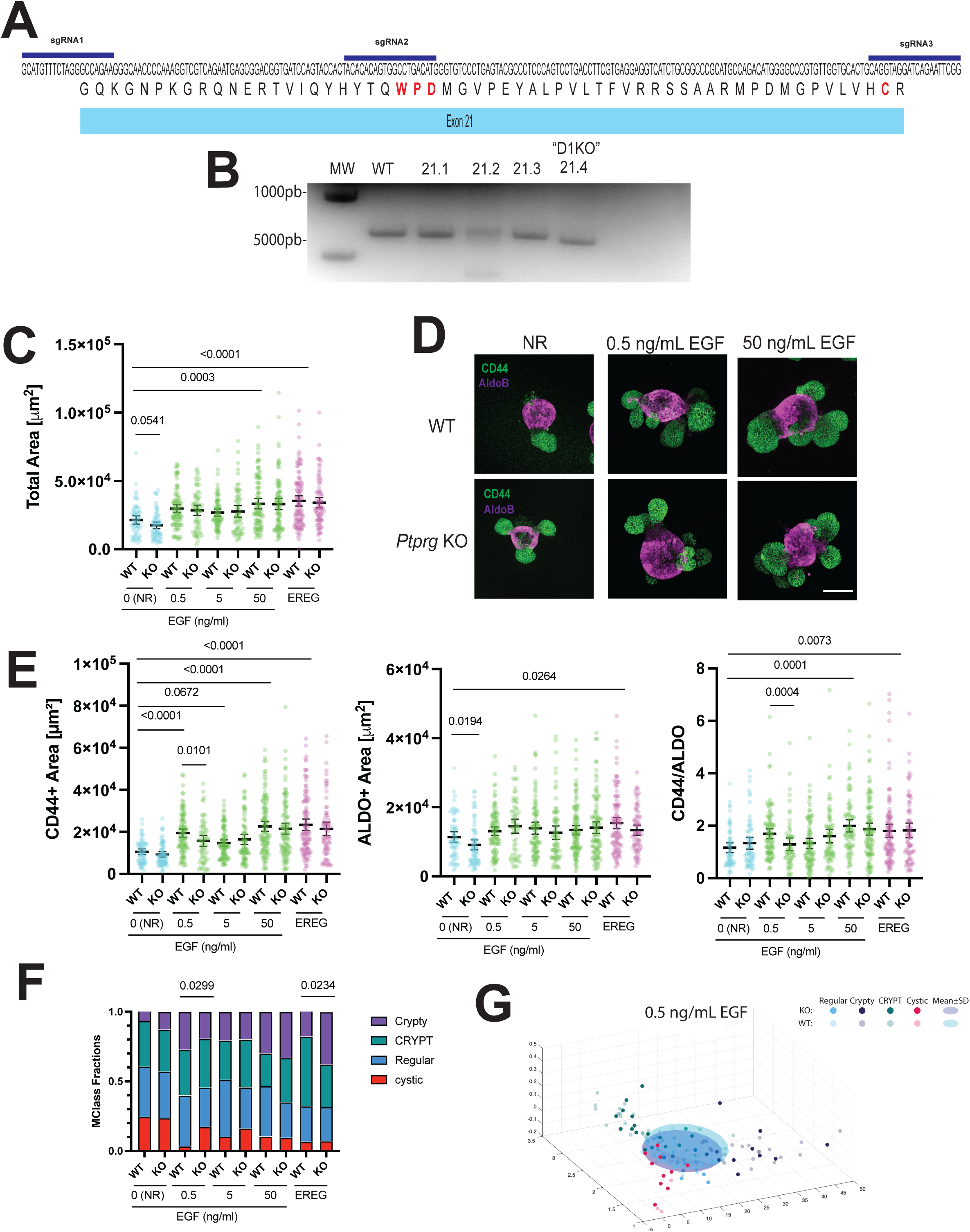
Sub-saturating EGF and EREG modify crypt growth and morphology in Ptprg KO organoids. (A) Genomic sequence and protein sequence of Ptprg Exon 21 showing the position of gRNAs used in this study. (B) Genotyping of WT and independent edited organoids (WT and Clones 1-4) using primers spanning the full length of Exon 21. Clone 21.4 or D1KO was used for further experiments (C-G). Single crypts were seeded and grown for 96 h with media containing 0 (NR media), 0.5, 5, and 50 ng/mL of EGF or 50 ng/mL of EREG. (C) Scatter plots showing mean ± 95% CI for total organoid area across all treatment conditions. Results obtained from n=66- 96 organoids in N=3 experiments. Significance between WT and KO organoids was calculated using Mann-Whitney’s test. Group comparisons to NR significance was calculated using ANOVA with Tukey’s test. p-values <0.05 are shown. (D) Representative images of mSIOs showing tissue transformation in WT and KO organoids; 50 ng/mL EGF organoids are shown as controls. Organoids were immuno- stained with CD44 (Green) and ALDOB (Magenta). Scale bar: 100 μm. (E) Scatter plots showing mean ± 95% CI of CD44, ALDOB areas, and the CD44/ALDOB ratio in all treatment conditions. Results obtained from n=66-96 organoids in N=3 experiments. Significance between WT and KO organoids was calculated using Mann- Whitney’s test. Group comparisons to NR were calculated using ANOVA with Tukey’s test. p-values <0.05 are shown. (F) Classification of organoid morphologies obtained from n=66-96 organoids in N=3 experiments as described above. Stacked bar plots of morphological Class (MClass) distribution. ‘Regular’ (blue) represents the prototypical crypt-villus structure; versus the aberrant classes: ‘Cystic’ (red) featureless, round organoids; ‘CRYPT’ (green) of enlarged crypts and ‘Crypty’ (violet) with enrichment of crypts. P-values: Chi^2^-test, p-values <0.05 are shown. (G) Representation of organoid morphologies in the morphometric space (principal components PC1, PC2, PC3). Representative PC of significantly altered morphologies, more examples in Figure S4 D1KO organoids (KO) and the mean±SD ellipsoid (solid colors; purple) versus wildtype (WT) and the mean±SD ellipsoid (pastel colors; aqua), visualizing the shift of morphological class distribution towards reduced crypt formation (‘Cystic’, red) in organoids harboring a Ptprg KO 0.5 ng/mL EGF media condition.

3 single guide RNAs (sgRNAs) targeting the 5’ and 3’ ends of exon 21 and the WPD encoding region were electroporated along with purified Cas9-NLS protein into a single-cell solution of mouse intestinal organoids (Figure S2A, B). We genotyped single clonal organoids with primers spanning exon 21; wt amplicons were predicted to be 660 bp, and edited alleles were expected to be shorter. Three (c2, c4, c5) out of the 12 clones tested showed different amplicons compared to wt (Figure 3B, Figure S2B). Because editing in c2 and c5 potentially yields 2 alleles with different outcomes, we selected c4 for sequencing (Figure S2B). Sequencing of c4 showed an inversion of exon 21 and the initial segment of the contiguous intronic region, alongside a deletion in the middle region of the exon where the WPD residues were located. (Figure S2C and D). This large-scale genetic modification is predicted to generate an early stop codon and completely disrupt the coding region of exon 21, likely generating a LoF mutation.

Since PTPRG shows low expression in intestinal tissue, where its expression is enriched in rare enteroendocrine cells (Human Protein Atlas, (Vezzalini et al., 2007)), we used nanoLC-MS/MS based proteomics to confirm PTPRG reduction in D1KO organoids. To improve PTPRG detection, we ran SDS-PAGE before tryptic digestion (Figure S3A). Full-length PTPRG is an ≈180 kDa protein with proteolytic cleavage products found at 120kDa or 100kDa; additionally, PTPRG is known to generate dimers and/or oligomers (Barr et al., 2009) which are often observed in Western blot as higher molecular weight bands in lysates from high PTPRG-expressing tissues (Moratti et al., 2015). Using Coomassie staining, we sliced the gel into 4 blocks between 250 and 100 kDa; we also sliced a block from a high-intensity band above the 250 kDa marker (Figure S3A, KM2813_DIA2_ID.xlsx). We consistently detected PTPRG in the WT samples from the band above 250 kDa (Figure S3B, KM2813_DIA2 KM2813_DIA2_ID.xlsx). Notably, PTPRG was not detectable in D1KO (hereafter “*Ptprg* KO”) in triplicate experiments compared to WT organoids (Figure S3B, KM2939_DIA_Upper int.xlsx), confirming that CRISPR-Cas9 gene editing significantly reduced protein expression in edited organoids.

As in the synthetic systems, we observed no change in organoid area in *Ptprg* KO organoids compared with WT in the presence of 50 ng/mL EGF, further highlighting the requirement of PM EGFR in oncogenic transformation. Surprisingly, total area and ALDOB+ area were significantly reduced without EGF; however, the CD44/ALDO ratio was maintained, suggesting balanced crypt and villus development even without *Ptprg*. In contrast, 0.5 ng/ml EGF significantly reduced crypt area and shifted the CD44/ALDO ratio from a crypt-growth-favoring state to a more balanced one, suggesting that lower EGF concentrations differentially alter organoid development, potentially by engaging PM-located receptors with the endocytic machinery and endomembrane-associated phosphatases. Since downregulation of PTPs is often accompanied by increases in EGFR phosphorylation (Joshi et al., 2023; Lin et al., 2026),pEGFR and total EGFR expression in WT and KO organoids grown with NR and 0.5 ng/ml of EGF. We observed higher levels of total EGFR in NR-grown organoids; addition of EGF greatly reduced EGFR levels as previously described (Figure S3C and D) (Caracci et al., 2026). *Ptprg* KO did not significantly affect phosphorylation levels in either condition (Figure S3C and D), suggesting that *Ptprg* deficiency might not affect EGFR activation at tissue level compared to cultured cells monolayers.

Morphological classification showed a reduction in cystic organoids and an increase in enlarged crypt and multiple crypt organoids under EGF or EREG treatment in both WT and *Ptprg* KO organoids (Figure 3F and G, Figure S4C), resembling what was observed in the acute oncoprotein expression models. Interestingly, 0.5 ng/mL EGF significantly shifted organoid morphology in *Ptprg* KO organoids by increasing the fraction of “Cystic” organoids; in contrast, EREG promoted the development of multiple-crypt *Ptprg* KO organoids (Figure 3F and G, Figure S4C), suggesting a role for this phosphatase and potentially for ERBB family receptors in crypt morphology.

## Discussion

Cell signaling is often studied in isolation, where activation or inhibition of a protein of interest is examined for its effect on downstream components of a signaling cascade, highlighting how biochemical networks are modified inside the cell. However, cells do not live in isolation; Maturana and Varela referred to the intrinsic and extrinsic factors within a living system as interacting in a “cognitive” manner, generating a recurrent and bidirectional flow of information with the environment (Koseska and Bastiaens, 2017; Maturana and Varela, 2012). In *in vitro* culture systems, researchers commonly modify and control the environment’s biochemical composition, particularly the presence of growth factors. In the present work, we aimed to interrogate how intrinsic oncoprotein signaling or loss of tumor suppressor activity differs across different media formulations containing EGF-family ligands EGF and EREG.

In this “cognitive” dialog, organoids are indeed capable of modifying their environment by expressing and secreting different growth factors. For instance, Paneth cells in mouse intestinal crypts produce EGF and Wnt ligands that help maintain the stem cell niche (Sato et al., 2011a). More recently, RNA-seq of stem and transit-amplifying cells, and Paneth cells, confirmed that the intestinal crypt also expresses high levels of Tgfa, Hbegf, and *Btc* under EGF-grown conditions (Belevich et al., 2026). Finally, qRT-PCR of bulk organoids grown in NR or sub-saturating EGF showed increased expression of *Egf, Hbegf, Tgfa*, and *Areg* (Caracci et al., 2026). Wether this ligands are expressed in the crypt region or proliferating cells is noy yet stablished as EGF is produced by villus region differentiated cells in human intestinal tissue (Childs et al., 2023). *In vivo*, oncogenic KRAS increases EREG expression in non-small-cell lung cancer carcinoma (NSCLC) and is associated with unfavorable prognosis (Sunaga et al., 2013). EREG is expressed in APC^min^ mouse tumors; however, it is not required for tumor development (Lee et al., 2004). Endogenous expression of HBEGF and AREG is known to enhance regeneration in injured intestinal tissue (El-Assal et al., 2008; Shao and Sheng, 2010; Su et al., 2013). In contrast, EGF does not increase after epithelial tissue damage; however, NRG1, an EGF-family ligand that does not bind EGFR, increased and also promoted organoid proliferation *in vitro* (Jardé et al., 2020).

Disentangling the role of each independent ligand expressed in the intestine would require generating KO lines for each EGF-family ligand; KO mouse models of single EGF-family ligands show no gross abnormalities in gastrointestinal development (Abud et al., 2021). Single and compounded EGF-family ligands KOs have been systematically studied in MDCK cells, showing that most ligands play redundant roles when it comes to activation of ERK; however, low-affinity ligands such as EREG promote ERK activity propagation in neighboring cells more efficiently than high- affinity ligands (Deguchi et al., 2024; Lin et al., 2022). How this differential role of certain ligands shapes tissue development remains unclear and likely involves EGFR endocytosis machinery and endosome engagement with organelle microdomains (Jendrisek et al., 2026 preprint). For mSIOs, we have argued that low- or high-affinity ligand regulation of the EGFR endocytic cycle balances proliferation and differentiation in crypt and villus regions, respectively (Caracci et al., 2026).

In the present work, we found that distinct media formulations containing (or lacking) EGF or EREG promote oncogenic tissue transformation by differentially modulating the proliferation-differentiation balance and/or by altering organoid morphology. Our chemically inducible protein stabilization methodology for oncoprotein activation is intended to illustrate early events in tissue transformation; whether long-term expression (>96 h) of these proteins can induce other phenotypes or generate *de novo* oncogenic mutations within the developing organoid remains to be elucidated. Without chemical activation with TMP or TMP-Cl, our synthetic constructs did not affect overall organoid morphology, whereas chronic expression of both KRASG12V and β-Ctn4A led to massive tissue transformation (Munoz-Nava et al., 2026 preprint; Soetje et al., 2026 preprint).

Organoids grown in the well-described basal media containing 50 ng/mL EGF showed no change in overall crypt or villus size in either synthetic or KO models; organoids grown in NR media (no added EGF) showed increased crypt growth in synthetic models and decreased growth in KO models. NR-grown organoids show high levels of EGFR at the plasma membrane, whereas organoids grown with ligands show a fraction of the receptor in endosomes, preventing ligand binding and PM signaling. The decreased endosomal activity in turn increases EGFR phosphorylation by promoting monomeric trans-phosphorylation and reducing the interaction with PTPs associated with endomembrane compartments (Baumdick et al., 2015; Stanoev et al., 2018).

Overall, we hypothesize that EGFR presence at the PM, and therefore its activity, is readily regulated by the affinity and concentration of EGF-family ligands in the media. Thus, saturating EGF concentrations throughout organoid development will continuously deplete receptors from the surface and send them to lysosomal degradation. In contrast, lower EGF concentrations or lower-affinity ligands such as EREG promote receptor recycling, maintaining the organoids’ ability to sense ligands produced by the epithelia or added exogenously to the culture media.

Morphometric analysis of organoids also showed a clear trend suggesting that addition of EGF or EREG promotes crypt fission in all conditions tested relative to NR-grown organoids. Crypt formation usually occurs about 48 h after single-crypt seeding and requires Paneth cells, which direct the first symmetry-breaking event in developing organoids toward a crypt/villus morphology (Serra et al., 2019). We have previously shown that EGF or EREG promote crypt growth in organoids; however, neither of them changes the number of Paneth cells (Caracci et al., 2026), suggesting that ligand activity may trigger acto-myosin-mediated apical constriction that precedes crypt formation in developing organoids (Hartl et al., 2019). Maintenance of Paneth cell numbers also suggests that EGF or EREG does not alter the number of stem cells within the niche, despite their effects on overall crypt growth. Crypt size results from cell proliferation and luminal inflation, which stretches cells within the organoid; luminal inflation also directly correlates with crypt fission rate in intestinal organoids (Tallapragada et al., 2021). We observe an increase in the percentage of organoids with enlarged crypts in TMP-treated β-Ctn4A-DD organoids at 50 and 5 ng/mL EGF, and although β-catenin and APC deficiency are known to enhance proliferation (Andreu et al., 2005; Hatzis et al., 2008), we cannot distinguish whether this enlargement is a result of proliferation or inflation with our methodology. Interestingly, activation of the mechanosensitive calcium channel Piezo1 has been shown to reduce crypt fission in organoids (Tallapragada et al., 2021). Notably, EGFR undergoes clathrin-mediated endocytosis in response to mechanical stress and Piezo1 activation in HeLa cells (Pardo-Pastor and Rosenblatt, 2023), suggesting a link between EGFR endocytic cycle and crypt fission events.

Notably, for *Ptprg* KO and β-Ctn4A-DD organoids, we observe that 0.5-5 ng/ml of EGF in the media induced changes in organoid development distinctive from NR media, affecting both crypt and villus development. On the other hand, the morphometric analysis of crypt organization showed that increased proliferation does not necessarily correlate with changes in the spatial organization of crypts, further highlighting that the establishment of crypts does not rely exclusively on proliferation rates.

Spatial proximity proteomics showed that EGFR and β-catenin are in close proximity early and late after EGF treatment; furthermore, EGF and EGFR seem to be required for expression of β-catenin target genes such as *cMYC* and *AXIN2* (Lin et al., 2026) Although this dynamical regulation seems to be context-dependent (Arasada et al., 2018; Knight et al., 2019). β-catenin can also be phosphorylated by EGFR, dissociating from E-cadherin complexes at the PM, increasing protein stabilization and transcriptional activity (Lilien and Balsamo, 2005). Thus, it is likely that early oncogenic β-catenin activity relies on the presence of EGFR at the PM, which, under basal media conditions is greatly reduced due to endocytic routing to degradation.

Expression levels of EGFR are tightly regulated, Lrig1 enhances ligand-dependent ubiquitination of EGFR (Laederich et al., 2004) and *Lrig1* KO leads to enlarged crypts in both colon and small intestine; furthermore, small intestinal organoids grown without EGF grow larger than controls (Wong et al., 2012). More recently, introduction point mutations in FBXW7, an E3 ubiquitin ligase commonly mutated in CRC, lead to increased levels of EGFR in human colon organoids. Edited organoids in turn showed higher cell surface expression of EGFR compared to WT, which allowed these organoids to grow normally with 0.005 ng/ul of EGF in the media, a 10,000-fold decreased compared to the amount required by control organoids (Boretto et al., 2024). Although phosphorylation related signaling activity of EGFR and it’s expression levels are certainly coupled, little is known of how EGFR phosphorylation regulates intestinal development. In these contexts, several protein tyrosine phosphatases are enriched in CRC (Wang et al., 2004) and also target EGFR phosphorylation (Stanoev et al., 2018). In addition, PTPRG downregulation has been documented in colorectal cancer cell lines and has further been shown to act as a tumour suppressor in breast cancer, nasopharyngeal carcinoma and chronic myeloid leukemia (Cheung et al., 2015; Della Peruta et al., 2010; Duan et al., 2025; Tomasello et al., 2020). Also, PTPRG expression is up-regulated by inflammation processes in the brain and liver (Brenachot et al., 2017; Lorenzetto et al., 2014). *PTPRG* KO MCF7 cells showed a dramatic increase in EGFR phosphorylation even in the absence of the activating ligand EGF, enhancing pro-migratory behavior and reducing proliferation(Joshi et al., 2023; Stanoev et al., 2018). Thus, generating a *Ptprg* KO organoid potentially offered the possibility to study the role of a constitutively active, oncogenic EGFR. Interestingly, in our *Ptprg* KO organoids, we found that without media EGF, organoids showed reduced villus and total area, although we could not detect any differences in EGFR phosphorylation by immunoblot, likely because PTPRG expression is potentially heterogeneous across multiple intestinal lineages, which has not been described. Immunohistochemistry of fixed tissue showed that PTPRG is sparsely expressed, with high levels in distinct, rare endocrine cells (Vezzalini et al., 2007). As described above, we hypothesize that EGFR maintenance at the PM is essential for observing PTPRG phenotypic activity during organoid development.

The link between EGFR and oncogenic KRAS has been widely documented, as pharmacological EGFR inhibitor efficiency is highly dependent on KRAS activity (Stanland et al., 2025). As for disease models, control over oncogenic KRASG12V expression and activity is fundamental, as overexpression leads to a hyper- proliferative spheroid, resembling other established cancer model organoids in both humans and mice (Matano et al., 2015). However, KRasG12V endogenous expression does not affect intestinal development in mice or Drosophila (Guerra et al, 2003, Vega-Cuesta et al, 2025), suggesting that expression levels are highly relevant for oncogene activity. In order to avoid overexpression, doxycycline-induced KRASG12V expression was able to induce pERK in mSIOs compared to controls; however, it did not modify organoid morphology (Brandt et al., 2019). In our model, KRASG12V oncoprotein signaling activity is modulated by control over the protein’s expression level and its sub-cellular localization, membrane and this approach has been successfully characterized in both organoids and MDCK cells (Munoz-Nava et al., 2026 preprint).

It is likely that associated of KRASG12V with EGFR is required for efficient interaction with downstream effectors either in the PM or endosomes, interaction that is greatly reduced due to saturating levels of EGF driving receptor degradation throughout a 96h developmental window (Caracci et al., 2026). Isogenic cell lines expressing KRASG12V showed that while KRAS activity is enhanced in the mutant isoform, the effect on downstream components such as pAKT and pERK is observed only after EGF treatment with no basal activation (Hood et al., 2019), suggesting synergistic effects between EGFR and RAS activity. No co-localization between Ras and EGFR has been observed within endosomal compartments after EGF stimulation (Surve et al., 2021), however Grb2 maintains EGFR co-localization within the endosomes (Fortian and Sorkin, 2014). In all, this idea requires further inquiry as association of EGFR with Ras proteins appears highly dependent on expression levels in cultured cells and a comprehensive revision of endosomal dynamics in organoids in currently not available.

Targeted expression of RasG12V in *Drosophila* midgut stem cells increased proliferation to similar levels as EGFR ligand overexpression, and both Ras and EGFR expression were required for proliferation (Jiang et al., 2011). KRASG12D increases EGFR expression and phosphorylation; furthermore, EGFR is necessary for tumorigenesis in pancreatic ductal adenocarcinoma (Ardito et al., 2012). Notably, EREG treatment greatly enhanced crypt growth in TMP-Cl-treated organoids; as mentioned above, EREG is also commonly upregulated in lung and pancreatic cancer. EREG also generates more sustained EGFR signaling by promoting receptor endocytosis and recycling toward the PM (Freed et al., 2017). EREG increased both crypt and villus areas in TMP-Cl-treated organoids, while EGF had no effect at any tested concentration, suggesting that EREG’s sustained signaling and receptor recycling promote oncogenic tissue transformation in response to KRAS G12V signaling activity.

Altogether, our results highlight how EGF-family growth factors commonly found in most organoid systems interact with oncogenic tumor suppressor mutations or acute activation of oncogenic protein signaling. Our observations and conclusions likely extend beyond intestinal organoids and cancer to other genetic perturbations in living systems aimed at modeling disease phenotypes. Saturating amounts of EGF or R- spondin have recently been used to deplete the PM of mSIOs of EGFR and Lgr5, respectively, to study secretory membrane trafficking from the Golgi (Belevich et al., 2026), further highlighting the impact of high concentrations of growth factors in the culture media.

## Methods

### Plasmids

All cDNAs were cloned into a piggyBac-CAG-IRES-PuroR vector (Wang et al., 2008) using the Gibson 475 Assembly Master Mix (NEB, cat. # E2611S) following the manufacturer’s protocol. Plasmids for β-catenin expression systems: pPB-bCtCDD- PCh-Hal and pPB-bCtC have been previously described in (Soetje et al., 2026, preprint) and plasmids for KRAS G12V expression: pPB-DDGDomG12V-THtKRAS and pPB-mKate2-KRASG12V have been described in (Munoz-Nava et al., 2026, preprint).

### Organoid Culture

Cryopreserved Mouse small intestinal organoids were purchased from STEMCELL Technologies (Cat #70931). Single crypts were expanded 30 μL Matrigel (Corning® Matrigel® Growth Factor Reduced (GFR) Basement Membrane Matrix, LDEV-Free, 354230) droplets in 24 well plates in ENR medium (Advanced Dulbecco’s modified Eagle’s medium/F12 with 100 u/mL penicillin/streptomycin, 10 mM HEPES, 2 mM Glutamax, 1x B27 (Life Technologies), 50 ng/mL mEGF (Peprotech, 315-09), 500 ng/mL R-Spondin 1 (Peprotech, 315-32), and 100 ng/mL Noggin (Peprotech, 250-38) and 1 mM N-acetylcysteine). Media was replaced with fresh media every 48h and organoid were expanded for 96-120 h after seeding.

EGF and EREG treatments.

For treatment medium without EGF, henceforth “NR medium” was supplemented with Epiregulin 50 ng/mL of EREG (Biotechne, 1195-EP-025) or decreasing concentrations of EGF (50, 5, 0.5 ng/ml). For oncoprotein stabilization of βCtn4A-DD, 10 µM of TMP (Trimethoprim T7883; Sigma-Aldrich) was added at seeding and replenished every 48 h. For oncoprotein Stabilization of CID-DD-KrasG12V, 1 µM TMP-Cl was added (ChiroBlock GmbH; Bitterfeld-Wolfen, Germany) at seeding and replenished every 48 h. After 96 h of treatment, organoids were fixed and immunostained for analysis.

### Immunoblotting

Organoid pellets were resuspended in modified RIPA buffer (50 mM Tris HCl, pH 7.4, 150 mM NaCl, 1 mM EDTA, 1 mM EGTA, 0.5% Triton X-100, and 0.5% sodium deoxycholate) supplemented with Complete Mini EDTA-free protease inhibitor (Roche Applied Science) and phosphatase inhibitor cocktail 2 and 3 (1:100, P5726 and P0044; Sigma-Aldrich), frozen in liquid nitrogen and stored in −70°C freezer. Samples were then mechanically dissociated using 10-gauge need syringes and centrifuged at 15,200g for 15 min at 4°C. Protein quantification was determined by BCA as described by manufacturer (23235; Thermo Fisher Scientific). 40 μg of protein was loaded in 8% SDS/PAGE gels and then transferred onto PVDF membranes. Membranes were then blocked in intercept buffer and incubated with primary antibody overnight. EGFR (C75B9) (1:1,000, 2646; Cell Signaling Technology) for total EGFR, EGFR Y1068 (AP774Y) (1:500; 40815; Abcam) for phospho-EGFR (pEGFR), and α-Tubulin (1:10,000; T6074; Merck) were used. Secondary antibodies IRDye 680 donkey anti-Rabbit IgG (1:10,000), IRDye 800 donkey anti-Mouse IgG (1:10,000) and IRDye 800 donkey anti-Rabbit IgG (1:10,000) (LI-COR Biosciences) were used. Membranes were documented using using Odissey CLx Imager and band intensity of full length EGFR, pEGFR and Tubulin was quantified using Fiji/ImageJ.

### Immunostaining

Single Crypts grown in 8-well labteks for 96 h under the described culture media treatments were fixed with 4% Paraformaldehyde solution in PBS for 40 min. After 3 washes with IF-buffer (0.2% Triton X100, 0.05% Tween20) for 5 min organoids were permeabilized with PBS 0.5% Triton X100 for 1h. Blocking was performed with IF buffer supplemented with 3% BSA. Primary antibodies CD44 (1:500, 560568, BD Pharmigen™) and Aldolase (1:500, ab153828, Abcam), were incubated overnight with agitation at 4°C. Organoids were then washed three times with IF Buffer for 5 min and then incubated for 1 h with Alexa Fluor secondary antibodies (1:500, Thermo Fisher Scientific) and DAPI (1:1,000). Alexa Fluor 488 (A11008, A11001), Alexa Fluor 546 (A11081, A11030), and Alexa Fluor 647 (A31573, A21247) secondary antibodies were used. After washing 3 times with IF buffer for 10 min, samples were kept in Clearing Solution (50% vol/vol Glycerol, 2.5 M Fructose) up to imaging at 4°C.

### Confocal Laser Scanning Microscopy (CLSM)

Organoids were imaged using Leica Confocal TCS SP8 operated with LASX software (Leica Microsystems) with a 20×/0.75 NA air objective equipped with Laser Kit WLL2 + Pulse Picker and 405 diode laser for illumination. 2μm z-stacks were acquired in all channels and area quantifications were performed in Max Intensity Projection binary masks of either DAPI, CD44 or AldoB representing total, crypt and villus area respectively. Image analyses were performed with Fiji/ImageJ software.

Quantification of β-Ctn4A-DD fluorescence intensity.

β-Ctn4A-DD mSIOs were seeded as single crypts in a 24-well glass bottom plate (Cellvis, cat. # P24-1.5H-N), and incubated in ENR medium at 37°C for 96 h with in ENR (Vehicle) or TMP (10 μM). mSIOs were imaged using an Olympus IX81 inverted microscope (Olympus Life Science) with MT20 illumination system and an incubation chamber at 37 °C and 5% CO_2_ atmosphere. Using a 10x/0.4 NA air objective and an ORCA-ER CCD digital camera (Hamamatsu Photonics, Japan). z-stacks of 120 µm with 6 slices were acquired in Transmission, mCitrine (YFP filter set) and mCherry (RFP filter set) every 2 h for up to 120 h organoid lifetime.

β-Ctn4A-Citrine intensity from β-Ctn4A-DD mSIOs was quantified using ImageJ/Fiji. From the mSIO z-stacks, maximum intensity projections were generated. Segmented binary mSIO masks were used to generate the corresponding organoid contour. Mean intensities of the peripheral cell layer were acquired by shrinking the mSIO contour by 10px and converting the area selection to a line selection with line thickness of 10px. This line selection covers the peripheral cell layer. β-Ctn4A-DD intensity in TMP or treated mSIOs was related to the mean intensity of the corresponding Vehicle controls.

Quantification of CID-DD-KRASG12V fluorescence intensity.

CID-DD-KRASG12V mSIOs were seeded from single crypts in a 24-well glass bottom plate (Cellvis, cat. # P24-1.5H-N), and incubated in ENR medium at 37°C for 96 h with DMSO or TMP-Cl (1 μM). We used CLSM, with a 20×/0.75 NA air objective, for imaging a representative focal plane from individual mSIOs. For quantification, we segmented CD-DD-GDom and TH-tKRAS pixels using local thresholding and extracted the mean for each organoid. Image processing and quantification were carried out using FIJI (fiji.sc) and the Python 3.11 (python.org) packages: imageio (pypi.org/project/ImageIO/), NumPy (numpy.org), pandas (pandas.pydata.org), scikit- image (scikit-image.org), SciPy (pypi.org/project/scipy/).

### Morphometric analysis

The morphometric analysis is described in detail in (Soetje et al., 2026, preprint). Here, we adopted the established analysis by adding two additional morphological features, resulting in a feature set of: Number of Crypt-like Features / Area-fraction of Largest Inscribed Circle, Relative Peak Width, Relative Peak Prominence, Aspect Ratio, Hu Moment 1, Hu Moment 2, Hu Moment 3, Hu Moment 4, Zernicke Moments Z20 and Z22. This set was selected from 41 morphometric features by minimum redundancy maximum relevance and univariate feature ranking, as well as predictor importance and removal or correlating features comparing Vehicle vs. Treatment/KO for all organoid models. A Principal component analysis was applied to generate and visualize a morphometric space.

The prototypical crypt-villus structure of normally developed mSIOs was defined by the mean+/- 0.7SD volume of PC1-PC3 from the respective ENR control / organoid model (β-Ctn4A-DD ENR, CID-DD-KRASG12V ENR and WT ENR). All organoids from other conditions within this volume were labeled as ‘control-like’. All non ‘control- like’ mSIO morphologies were clustered via k-means clustering and the respective aberrant clusteres assigned to the ‘Crypty’ crypt enriched, ‘CRYPT’ crypt enlarged, or ‘Cystic’ featureless morphological groups, as visible from the localization of that cluster in the morphometric space where PC1 is dominated by the feature umber of Crypt-like Features / Area-fraction of Largest Inscribed Circle, and PC2 by the AR while PC3 is incorporating rel. crypt dimensions and asymmetry from the Zernicke moment Z22. The core organization of that group as again defined by mean+/- 0.7SD and used for classifier training, generating one classifier for each organoid model used.

### Electroporation

mSIOs were transfected by electroporation using the NEPA21 electroporator 545 (Nepagene, Japan) as previously described (Fujii et al., 2015). WT organoids were grown for 48 h in ENR media before adding 1.25% v/v DMSO, CHIR-99021 (Sigma- Aldrich), and Y-27632 (Holzel Biotech, cat. # M1817-50 mg) to ENR medium without Penicillin/Streptomycin for 48 h. After treatment, 6 wells of mouse small intestinal organoids grown in 24-well plates were pooled and digested with Trypsin LE (Gibco, cat. # 12605010) containing Y27632 for 7 minutes before vortexing and then for another 7 minutes. Complete digestion into single cells was monitored by brightfield microscopy. Cells were then pelleted and washed once with Opti-MEM (Gibco, cat. #31985070), media before electroporation. For the generation of stable organoid lines 10g of Plasmid DNA and 3 g piggyBac transposase vector were added to 100 L of Opti-MEM and used to resuspend the single-cell organoid pellet. The cell suspension with Plasmid mixture was added into a 2 mm gap cuvette (Xceltis, EC-002S) for electroporation. Poring/transfer pulse 555 conditions were used as previously described (Fujii et al, 2015): Voltage – 175/20 V, 556 Pulse length – 5/50 ms, Pulse interval – 50/50 ms, Number of pulses – 2/5, Decay rate 557 10/40 %, Polarity +/+/-. After electroporation, the cell suspension was resuspended in warm ENR medium without Penicillin/Streptomycin for 30 min, centrifuged, and seeded in Matrigel domes. Puromycin (3 µg/ml) was added 96h after electroporation for selection of stably transfected organoids.

### RNP CRISPR-Cas9 editing

For CRISPR-Cas9 editing, organoids were pre-treated as described above for electroporation. sgRNAS and spCas9 2NLS were purchased from Synthego (Redwood, CA). sgRNAs were designed using the Synthego Knockout Guide Design tool (https://design.synthego.com/#/). 3 independent sgRNAs encompassing the full length of *Ptprg* exon 21 were designed (sgRNA1: GCATGTTTCTAGGGCCAGAA; sgRNA2:ATGTCAGGCCACTGTGTGTA; sgRNA3: CAGGTAGGATCAGAATTCGG).

50 pmol of each sgRNA resuspended in TE buffer were mixed with 30 pmol of Cas9 2 NLS (5:1 ratio) in 10 µL of OptiMEM and incubated at RT for 10 min. Afterward, 90 µL of OptiMEM was added to the mixture and used to resuspend the single-cell organoid pellet, then electroporated it as previously described using a Super Electroporator NEPA 21 Type II (Nepagene) (Fujii et al., 2015). Electroporated organoids were grown for 5-7 days before single organoids were isolated for clonal expansion. Single clones were genotyped by total DNA extraction using QuickExtract^TM^ DNA Extraction Solution (Biosearch^TM^ Techonlogies) and primers flanking the edited exon. Primers were designed to generate 660 bp amplicons for WT unedited alleles (Fw: GTCAGGAACGGGAGACCTC; Rv: ATAGTCAGTCAAACAGATGT). PCR was conducted using Q5® High-Fidelity 2X Master Mix (New England Biolabs), and amplicons were resolved in 2% Agarose Gels. Clones showing edited alleles were further expanded and sequenced. Amplicons were excised and purified from Agarose gels using Monarch® DNA Gel Extraction Kit (New England Biolabs) and cloned into plasmids using CloneJET PCR Cloning Kit (ThermoFisher Scientific). A minimum of 5 single bacterial colonies were expanded and purified using NucleoSpin® Plasmid (Macherey-Nagel) and sequenced using the provided sequencing primer. Obtained Clones were named according to the exon targeted and the clone number (e.g. Clone 21.2 corresponds to editing of exon 21 and the clone number 2).

### NanoLC-MS/MS proteomics for targeted PTPRG detection

Lysates were obtained as described in the “*Immunoblot*” section above. The lysates were reduced with β-ME in sample buffer and boiled for 5 min at 95°C and then with DTT added to 1mM for 30 min at RT. For alkylation, CAA was added to 5.5 mM final concentration for 20 min at RT. 40 μg of sample was run in a 6% gel SDS page gel casted in single-use 1,5 mm cassetes 40ug for 1h and 35 min at constant 30mA. Then gel was stained with Coomasie Brilliant Blue G (Sigma B-0770, MW 854.0) and then distained overnight with Methanol/Acetic Acid solution.

Resulting protein bands were sliced from the gel using a sterile scalpel into small pieces (ca. 1x1 mm) and put into 2 ml Eppendorf tubes. For destaining 3 washes with 2 ml 50 mM ABC( 20mM NH_4_HCO_3_) + 50 % EtOH for 15 min at 30°C with 1000 rpm agitation were used. For dehydration, the supernatant was incubated with 2 mL of ETOH for 10 min at 24°C with 1000 rpm agitation the EtOH was removed and excess was removed by 30 min centrifugation at 30°C in SpeedVac centrifuge (Eppendorf Concentrator Plus 5305 System). Trypsin stock solution (0,1 µg/µL in 10 mM HCl) (03708985001 from Sigma) was diluted 1:5 in Trypsin dilution buffer (100 mM Tris, 1 mM CaCl_2_, pH 8.5) to a 25 μL volume and added to the sample. After 15 min incubation at RT, 75 μL of 20 mM ABC was added and incubated overnight at 37°C with 1000 rpm agitation. Peptides were then extracted by adding TFA to 0.2% final concentration, incubated for 2min at 2000 rpm and then sequentially treated for 15 min at 24°C, 1000 rpm agitation with 30% ACN, 0.1% TFA; 60% ACN, 0,1% TFA;and 100% ACN. ACN was then evaporated using SpeedVac centriguge for 60 min at 30°C. Peptides were purified using StageTip filtration (AttractSPE^TM^ Disks Bio – C18, Catalog #SPE-Disks- Bio-C18-100.47.20, 20 SPE Disks, 47 mm, Lot: PP12BPO-5; Affinisep) and eluted with (60% ACN in 0.1% TFA in ddH_2_0) and excess was evaporated in SpeedVac centrifuge.

For relative protein quantification, tryptic peptides were separated and analyzed by nano-HPLC-MS/MS. The separations were carried out on UltiMateTM 3000 RSLCnano system (Thermo Scientific, Germany). The MS and MS/MS experiments were carried out on a Fusion Lumos™ Tribrid Orbitrap Mass Spectrometer equipped with a nano-spray source (Nanospray Flex Ion Source, Thermo Scientific). All solvents were LC-MS grade. The lyophilized tryptic peptides were dissolved in 20 µL 0.1% TFA in water. 3 µL of sample were injected onto a pre-column cartridge (5 µm, 100 Å, 300 µm ID * 5 mm, Thermo Scientific, Germany) using 0.1% TFA in water as eluent with a flow rate of 30 µL/min. Desalting was performed for 5 min with eluent flow to waste followed by back-flushing of the sample during the whole analysis from the pre-column to the PepMap100 RSLC C18 nano-HPLC column (2 µm, 100 Å, 75 µm ID × 50 cm, nanoViper, Dionex, Germany) using a linear gradient starting with 96% solvent A (water containing 0.1% formic acid) and 4% solvent B (acetonitrile containing 0.1% formic acid), increasing to 8% solvent B within 1 min, further increasing to 24.8% solvent B within 29 min and to 35.2% within 5 min and finally increasing to 90% solvent B within 2 min using a flow rate of 300 nL/min. The nano-HPLC was online coupled to the Mass Spectrometer using an un-coated SilicaTip (Tip-ID 10 μM, Fossiliontech, Spain). Mass range of m/z 350 to 1000 was acquired with a resolution of 120000 for full scan, followed by data-independent high energy collision dissociation (HCD) MS/MS scans with an isolation window of 18.6 m/z, a window overlap of 1 m/z and a loop time of 3 s. MS/MS spectra were measured with a resolution of 30000, a NCE of 30% and mass range mode set to auto. Data evaluation was performed using DIA-NN software (v.2.3.0 academia) (Demichev et al., 2020). The spectral library was predicted using a combination of a mouse database from uniprot and contamination database. The search was performed for full enzymatic trypsin cleavages allowing one miscleavage. For protein modifications carbamidomethylation was chosen as fixed modifications. The mass accuracy settings were optimized by DIA-NN. MBR was enabled and a cross-run retention time dependent normalization was performed. The false discovery rate was set to 1%. All experiments were performed in technical triplicates. Intensity values for PTPRG used from the pg_matrix.tsv table were plotted to compare PTPRG intensities in WT compared to knock-out samples (clone 21.4) using GraphpadPrism.

## Acknowledgements

We used Grammarly to assist with grammar, spelling, and stylistic improvements in the final draft of this work. Figure 1A and 2A were created using Biorender.com. We thank Andreas Brockmeyer, Sasikala Thavam and the Mass Spectrometry facility at Max Planck Institute of Molecular Physiology for technical assistance. Prof. Dr. PIH Bastiaens, passed away on May 15, 2025. All authors agree that his inclusion as a co- author is appropriate to honor his intellectual contribution to this work.

## Data and resource availability

Scripts and Data can be provided on reasonable request.

## Author Contributions

MO Caracci: conceptualization, data curation, formal analysis, investigation, visualization, methodology, and writing—original draft, review and editing., S Seidler: validation, methodology. LM Muñoz-Nava: conceptualization, investigation, data curation, formal analysis, visualization and editing. B Soetje: conceptualization, investigation, data curation, formal analysis, visualization and editing. K Michel: methodology. P. Janning: conceptualization, data curation formal analysis. PIH Bastiaens: conceptualization, funding acquisition, and project administration

---

**Figure S1.**
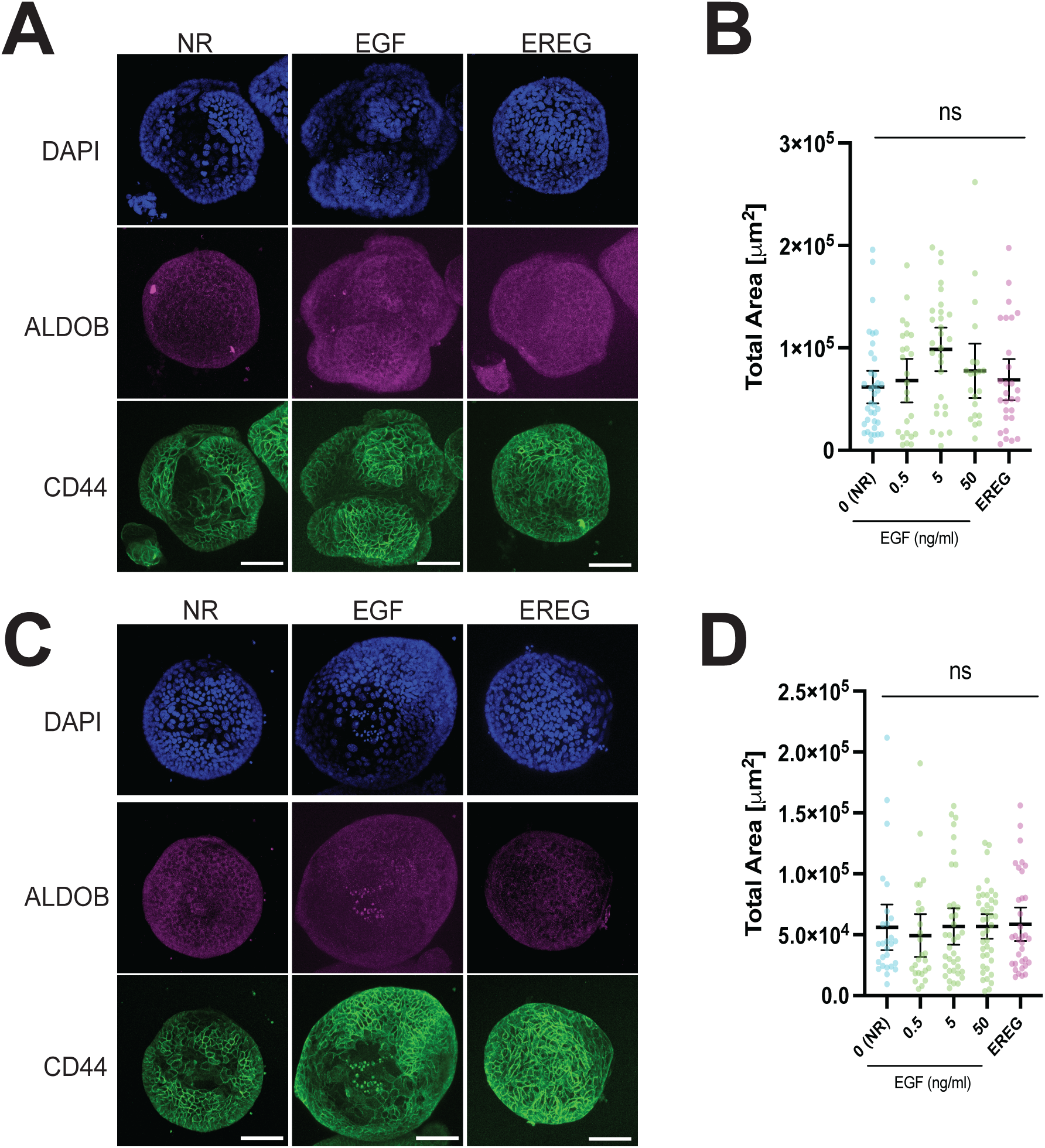
Constitutive expression of oncogenic KRASG12V and β-catenin4A disrupts crypt-villus organoid organization regardless media composition. (A- D) Single crypts were seeded and grown for 96h with media containing 0 (NR media), 0.5, 5, and 50 ng/mL of EGF or 50 ng/mL of EREG. (A) Representative images of mSIOs with constitutive β-catenin4A expression, grown with 0 ng/mL of EGF (NR), 50 ng/mL of EGF, or 50 ng/mL of EREG. Organoids were immuno-stained with CD44 (Green), AldoB (Magenta). Scale bar: 100 μm. (B) Scatter plots showing mean ± 95% CI of Total organoid area in all treatment conditions. Results obtained from n=26-40 organoids in N=3 experiments. Significance between treatments was calculated using Mann-Whitney’s test; ns, non-significant. (C) Representative images of mSIOs constitutively expressing KRASG12V and grown with 0 ng/mL EGF (NR), 50 ng/mL EGF, or 50 ng/mL EREG. Organoids were immuno-stained with CD44 (Green), AldoB (Magenta). Scale bar: 100 μm. (D) Scatter plots showing mean ± 95% CI of Total organoid area in all treatment conditions. Results obtained from n=21-36 organoids in N=3 experiments. Significance between treatments was calculated using Mann- Whitney’s test; ns, non-significant.

**Figure S2.**
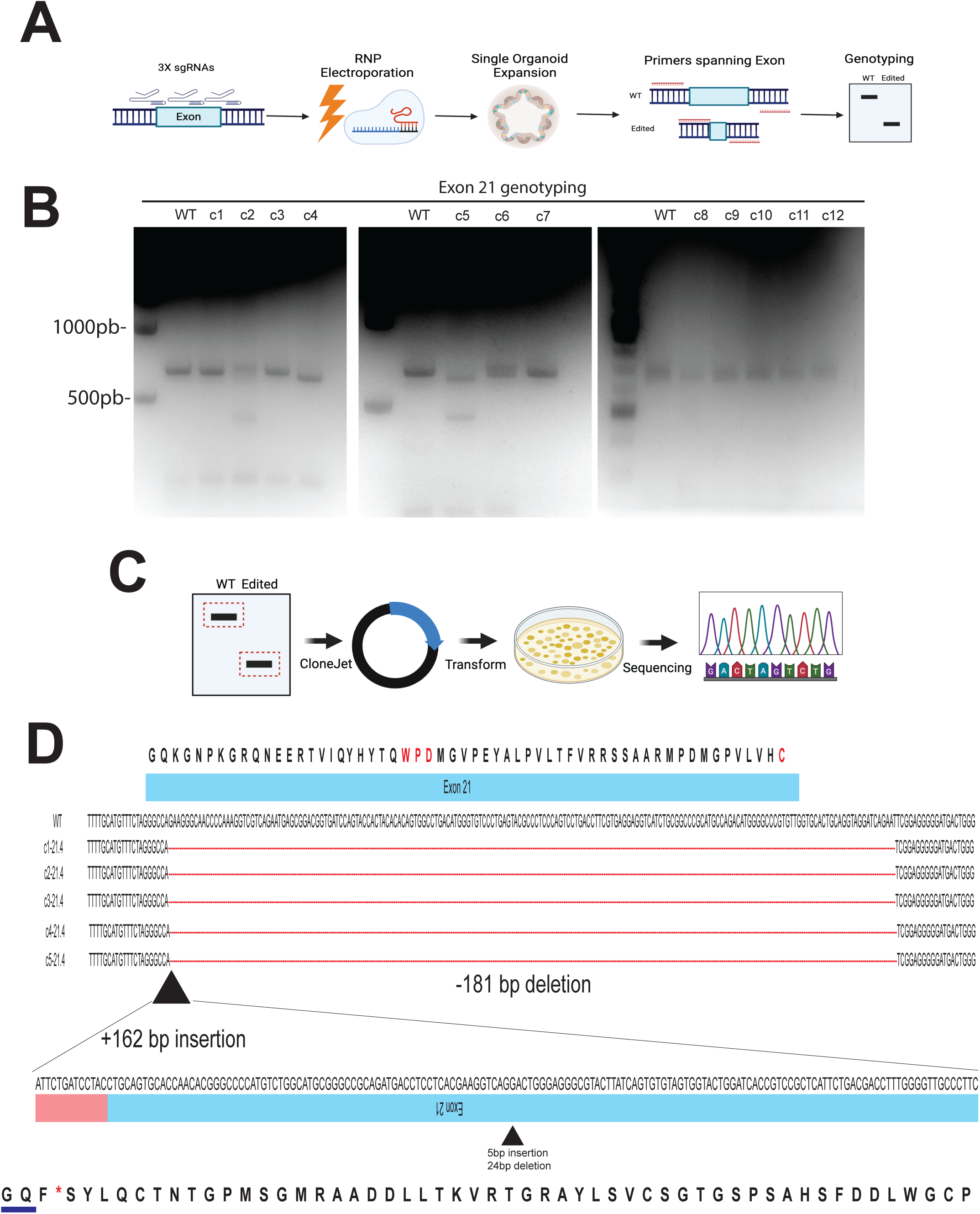
Gene editing of *Ptprg* Exon 21 leads to removal of catalytic residues of D1 phosphatase domain. (A) General strategy for CRISPR-Cas9 gene editing by electroporation of RNPs conjugated with 3 independent sgRNAs targeting exon 21. (B) Genotyping of WT and edited organoids (WT and Clones 1-12) using primers spanning the full length of Exon 21. (C) General strategy showing subcloning into pJet plasmids of PCR products obtained from genotyping in (C); the obtained plasmids were amplified in E. coli, purified, and sequenced. (D) Sequencing of genotyping products obtained from wt organoids and 21.4 clone, multiple plasmid clones result re shown for 21.4 (Clones 1-5), Red letters show critical residues for phosphatase activity located within Exon 21 in the D1 phosphatase domain. Red dotted line denotes a 181 bp deletion within edited organoid 21.4 clone. Dark triangle shows position of novel 162 bp insertion, corresponding to an inverted Exon 21 region. With the remaining first two codons of exon 21 (“GQ” aminoacids, blue underline) the resulting reading frame leads to an early STOP codon and removal of critical residues required for PTPRG phosphatase activity.

**Figure S3.**
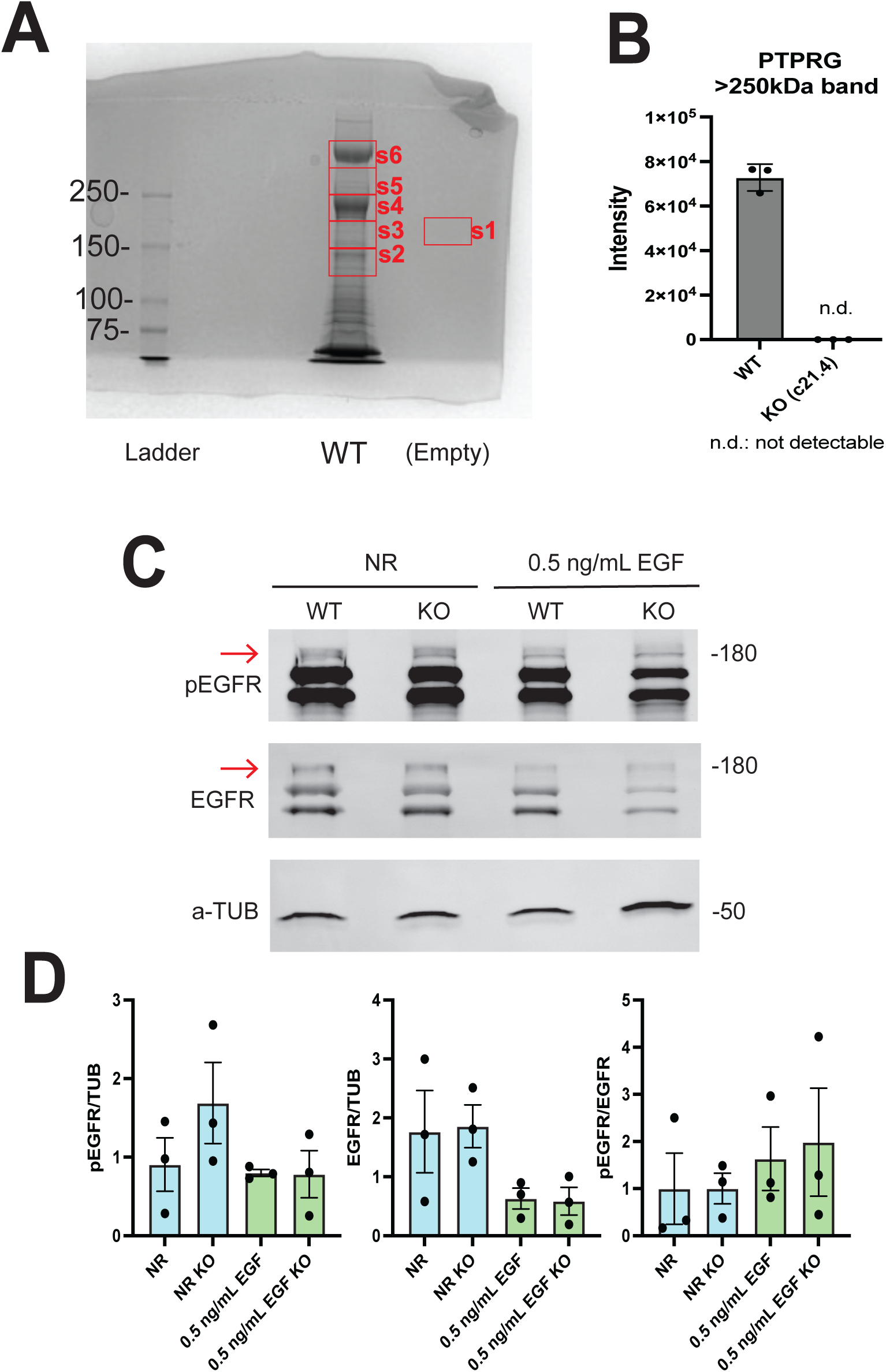
PTPRG is not detected in *Ptprg* KO organoids and EGFR phosphorylation is not affected in *Ptprg* KO organoids. (A) (Left) Total protein extracts from organoids grown for 96 h with normal basal medium (50 ng/mL EGF) were run on a 6% SDS-PAGE gel and stained with Coomassie. Based on expected MW of PTPRG full length protein, known isoforms and proteolytic by-products, bands between the 150 and above 250 kDa markers (s2-s6) were sliced and processed for nanoLC-MS/MS proteomics. A lane with no protein loaded (Empty) was used as a negative control (s1). (B) Bar Graph showing signal intensity of PTPRG protein detected by nanoLC-MS/MS proteomics in the high MW band. Results from 3 biological replicates of WT and KO protein extracts. Non-detectable n.d. (C) Immunoblot of WT and *Ptprg* KO organoids grown for 96 h in NR (0 ng/mL EGF) or 0.5 ng/mL EGF. Immunoblot of pEGFR, EGFR and a-TUB with approximate MW in kDa. Red arrows point to full-length EGFR or pEGFR used for band intensity measurements (D). Bar Graphs of the mean ± SEM of pEGFR or EGFR relative to a- TUB and the pEGFR/EGFR ratio. Results from 3 biological replicates, N=3.

**Figure S4.**
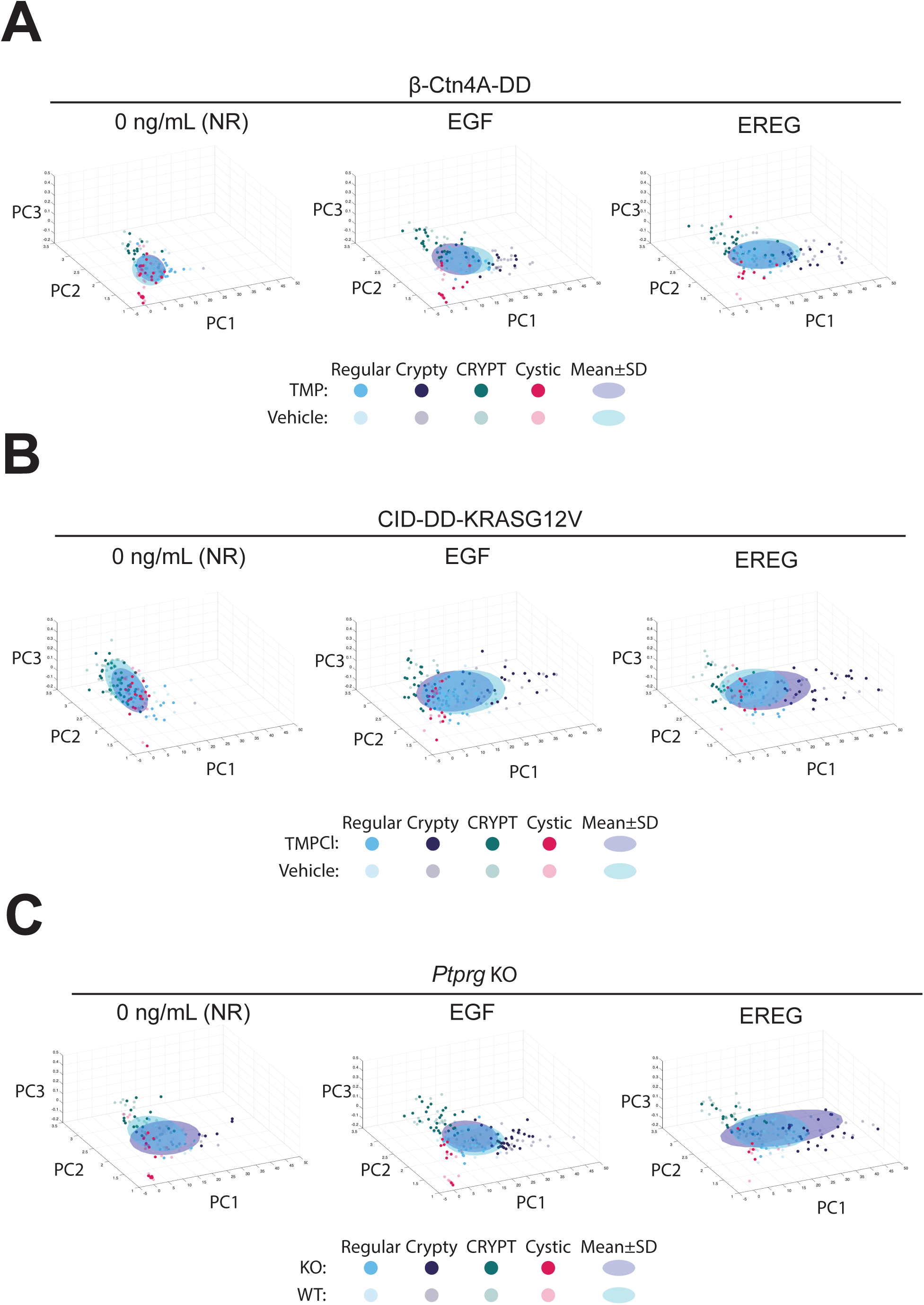
Morphological organoid distributions of NR, EGF and EREG conditions in synthetic bio-orthogonal systems and *Ptprg* KO. (A-C) Single crypts were seeded and grown for 96 h with media containing 0 (NR media), 50 ng/mL of EGF (EGF) or 50 ng/mL of EREG (EREG). Scatter plot of organoid morphologies in the morphometric space (principal components PC1, PC2, PC3). (A) β-Ctn4A-DD organoids grown in presence of TMP (TMP) and the mean±SD ellipsoid (solid colors; purple) versus vehicle-control (Vehicle) and the mean±SD ellipsoid (pastel colors; aqua). (B) CID- DD-KRASG12V organoids grown in presence of TMPCl (TMPCl) and the mean±SD ellipsoid (solid colors; purple) versus vehicle-control (Vehicle) and the mean±SD ellipsoid (pastel colors; aqua). (C) D1KO organoids (KO) and the mean±SD ellipsoid (solid colors; purple) versus wildtype organoids (WT) and the mean±SD ellipsoid (pastel colors; aqua).

## References

1. Abud, H. E., Chan, W. H. and Jardé, T. (2021). Source and impact of the EGF family of ligands on intestinal stem cells. Front. Cell Dev. Biol. 9, 685665.

2. Arasada, R. R., Shilo, K., Yamada, T., Zhang, J., Yano, S., Ghanem, R., Wang, W., Takeuchi, S., Fukuda, K. and Katakami, N. (2018). Notch3-dependent β- catenin signaling mediates EGFR TKI drug persistence in EGFR mutant NSCLC. Nat. Commun. 9, 3198.

3. Ardito, C. M., Grüner, B. M., Takeuchi, K. K., Lubeseder-Martellato, C., Teichmann, N., Mazur, P. K., DelGiorno, K. E., Carpenter, E. S., Halbrook, C. J. and Hall, J. C. (2012). EGF receptor is required for KRAS-induced pancreatic tumorigenesis. Cancer Cell 22, 304–317.

4. Barr, A. J., Ugochukwu, E., Lee, W. H., King, O. N. F., Filippakopoulos, P., Alfano, I., Savitsky, P., Burgess-Brown, N. A., Müller, S. and Knapp, S. (2009). Large-scale structural analysis of the classical human protein tyrosine phosphatome. Cell 136, 352–363.

5. Baumdick, M., Brüggemann, Y., Schmick, M., Xouri, G., Sabet, O., Davis, L., Chin, J. W. and Bastiaens, P. I. H. (2015). EGF-dependent re-routing of vesicular recycling switches spontaneous phosphorylation suppression to EGFR signaling. Elife 4, e12223.

6. Belair, D. G., Visconti, R. J., Hong, M., Marella, M., Peters, M. F., Scott, C. W. and Kolaja, K. L. (2020). Human ileal organoid model recapitulates clinical incidence of diarrhea associated with small molecule drugs. Toxicology in Vitro 68, 104928.

7. Belevich, I., Porcile, L., Sola-Carvajal, A., Grommisch, D., Annusvar, K., Heinz, P., Das, S., Webb, A. T., Andersson, S. and Pentinmikko, N. (2026). Golgi organization regulates stem cell function in the small intestine. Nat. Commun. 17, 7606.

8. Boni, C. and Sorio, C. (2022). The role of the tumor suppressor gene protein tyrosine phosphatase gamma in cancer. Front. Cell Dev. Biol. 9, 768969.

9. Boretto, M., Geurts, M. H., Gandhi, S., Ma, Z., Staliarova, N., Celotti, M., Lim, S., He, G.-W., Millen, R. and Driehuis, E. (2024). Epidermal growth factor receptor (EGFR) is a target of the tumor-suppressor E3 ligase FBXW7. Proceedings of the National Academy of Sciences 121, e2309902121.

10. Bos, J. L., Rehmann, H. and Wittinghofer, A. (2007). GEFs and GAPs: critical elements in the control of small G proteins. Cell 129, 865–877.

11. Brandt, R., Sell, T., Lüthen, M., Uhlitz, F., Klinger, B., Riemer, P., Giesecke- Thiel, C., Schulze, S., El-Shimy, I. A. and Kunkel, D. (2019). Cell type- dependent differential activation of ERK by oncogenic KRAS in colon cancer and intestinal epithelium. Nat. Commun. 10, 2919.

12. Brüggemann, Y., Karajannis, L. S., Stanoev, A., Stallaert, W. and Bastiaens, P. I. H. (2021). Growth factor–dependent ErbB vesicular dynamics couple receptor signaling to spatially and functionally distinct Erk pools. Sci. Signal. 14, eabd9943.

13. Caracci, M. O., Avila, M. E., Espinoza-Cavieres, F. A., López, H. R., Ugarte, G. D. and De Ferrari, G. V (2021). Wnt/β-catenin-dependent transcription in autism spectrum disorders. Front. Mol. Neurosci. 14, 764756.

14. Caracci, M. O., Seidler, S., Muñoz-Nava, L. M., Soetje, B., Michel, K. and Bastiaens, P. I. H. (2026). Proliferation and differentiation in intestinal organoids are balanced by ligand-modulated EGFR trafficking. Life Sci. Alliance 9, e202603733.

15. Childs, C. J., Holloway, E. M., Sweet, C. W., Tsai, Y.-H., Wu, A., Vallie, A., Eiken, M. K., Capeling, M. M., Zwick, R. K. and Palikuqi, B. (2023). EPIREGULIN creates a developmental niche for spatially organized human intestinal enteroids. JCI Insight 8,.

16. Clevers, H. and Nusse, R. (2012). Wnt/β-catenin signaling and disease. Cell 149, 1192–1205.

17. De Ferrari, G. V and Moon, R. T. (2006). The ups and downs of Wnt signaling in prevalent neurological disorders. Oncogene 25, 7545–7553.

18. Deguchi, E., Lin, S., Hirayama, D., Matsuda, K., Tanave, A., Sumiyama, K., Tsukiji, S., Otani, T., Furuse, M. and Sorkin, A. (2024). Low-affinity ligands of the epidermal growth factor receptor are long-range signal transmitters in collective cell migration of epithelial cells. Cell Rep. 43,.

19. Demichev, V., Messner, C. B., Vernardis, S. I., Lilley, K. S. and Ralser, M. (2020). DIA-NN: neural networks and interference correction enable deep proteome coverage in high throughput. Nat. Methods 17, 41–44.

20. Drosten, M. and Barbacid, M. (2020). Targeting the MAPK pathway in KRAS-driven tumors. Cancer Cell 37, 543–550.

21. Duan, H., Jia, J., Wang, T., Khan, G. J., Wang, L., Zhang, S., Yang, J., He, C., Li, F. and Hu, H. (2025). miR-23b is a negative regulator of tumor suppressing PTPRG in colorectal cancer. Transl. Oncol. 59, 102447.

22. El-Assal, O. N., Paddock, H., Marquez, A. and Besner, G. E. (2008). Heparin- binding epidermal growth factor–like growth factor gene disruption is associated with delayed intestinal restitution, impaired angiogenesis, and poor survival after intestinal ischemia in mice. J. Pediatr. Surg. 43, 1182–1190.

23. Fortian, A. and Sorkin, A. (2014). Live-cell fluorescence imaging reveals high stoichiometry of Grb2 binding to the EGF receptor sustained during endocytosis. J. Cell Sci. 127, 432–444.

24. Freed, D. M., Bessman, N. J., Kiyatkin, A., Salazar-Cavazos, E., Byrne, P. O., Moore, J. O., Valley, C. C., Ferguson, K. M., Leahy, D. J. and Lidke, D. S. (2017). EGFR ligands differentially stabilize receptor dimers to specify signaling kinetics. Cell 171, 683–695.

25. Fujii, M., Matano, M., Nanki, K. and Sato, T. (2015). Efficient genetic engineering of human intestinal organoids using electroporation. Nat. Protoc. 10, 1474– 1485.

26. Hart, M. J., De Los Santos, R., Albert, I. N., Rubinfeld, B. and Polakis, P. (1998). Downregulation of β-catenin by human Axin and its association with the APC tumor suppressor, β-catenin and GSK3β. Current Biology 8, 573–581.

27. Hood, F. E., Klinger, B., Newlaczyl, A. U., Sieber, A., Dorel, M., Oliver, S. P., Coulson, J. M., Blüthgen, N. and Prior, I. A. (2019). Isoform-specific Ras signaling is growth factor dependent. Mol. Biol. Cell 30, 1108–1117.

28. Iwamoto, M., Björklund, T., Lundberg, C., Kirik, D. and Wandless, T. J. (2010). A general chemical method to regulate protein stability in the mammalian central nervous system. Chem. Biol. 17, 981–988.

29. Jardé, T., Chan, W. H., Rossello, F. J., Kahlon, T. K., Theocharous, M., Arackal, T. K., Flores, T., Giraud, M., Richards, E. and Chan, E. (2020). Mesenchymal niche-derived neuregulin-1 drives intestinal stem cell proliferation and regeneration of damaged epithelium. Cell Stem Cell 27, 646–662.

30. Jendrisek, G., Mesa, D., Freddi, S., Miloro, G., Tordonato, C., Benvenuto, A. F., Quarto, M., Caputo, M., Raimondi, A. and Caldieri, G. (2026). Decoding EGFR ligand bias through an endocytic organelle platform. bioRxiv 2026–2028.

31. Jiang, H., Grenley, M. O., Bravo, M.-J., Blumhagen, R. Z. and Edgar, B. A. (2011). EGFR/Ras/MAPK signaling mediates adult midgut epithelial homeostasis and regeneration in Drosophila. Cell Stem Cell 8, 84–95.

32. Joshi, M. S., Stanoev, A., Huebinger, J., Soetje, B., Zorina, V., Roßmannek, L., Michel, K., Müller, S. A. H. and Bastiaens, P. I. H. (2023). The EGFR phosphatase RPTPγ is a redox-regulated suppressor of promigratory signaling. EMBO J. 42, e111806.

33. Klein, J. A., Heidmann, J. D., Kiyota, T., Fullerton, A., Homan, K. A. and Co, J. Y. (2025). The differentiation state of small intestinal organoid models influences prediction of drug-induced toxicity. Front. Cell Dev. Biol. 13, 1508820.

34. Knight, C., James, S., Kuntin, D., Fox, J., Newling, K., Hollings, S., Pennock, R. and Genever, P. (2019). Epidermal growth factor can signal via β-catenin to control proliferation of mesenchymal stem cells independently of canonical Wnt signalling. Cell. Signal. 53, 256–268.

35. Koseska, A. and Bastiaens, P. I. H. (2017). Cell signaling as a cognitive process. EMBO J. 36, 568–582.

36. Kotani, T., Ihara, N., Okamoto, S., Setiawan, J., Konno, T., Saito, Y., Murata, Y. and Matozaki, T. (2021). Role of Ras in regulation of intestinal epithelial cell homeostasis and crosstalk with Wnt signaling. PLoS One 16, e0256774.

37. Krishna, A., Meynert, A., Dolt, K. S., Kelder, M., Mesropian, A., Ewing, A., Brouwers, C., Claassens, J. W. C., Linssen, M. M. and Sheraz, S. (2026). Mutational scanning reveals oncogenic CTNNB1 mutations have diverse effects on signaling. Nat. Genet. 58, 366–375.

38. Laederich, M. B., Funes-Duran, M., Yen, L., Ingalla, E., Wu, X., Carraway, K. L. and Sweeney, C. (2004). The leucine-rich repeat protein LRIG1 is a negative regulator of ErbB family receptor tyrosine kinases. Journal of Biological Chemistry 279, 47050–47056.

39. Lee, D., Pearsall, R. S., Das, S., Dey, S. K., Godfrey, V. L. and Threadgill, D. W. (2004). Epiregulin is not essential for development of intestinal tumors but is required for protection from intestinal damage. Mol. Cell. Biol. 24, 8907–8916.

40. Lemmetyinen, T. T., Viitala, E. W., Wartiovaara, L., Kaprio, T., Hagström, J., Haglund, C., Katajisto, P., Wang, T. C., Domènech-Moreno, E. and Ollila, S. (2023). Fibroblast-derived EGF ligand neuregulin 1 induces fetal-like reprogramming of the intestinal epithelium without supporting tumorigenic growth. Dis. Model. Mech. 16, dmm049692.

41. Li, X., Nadauld, L., Ootani, A., Corney, D. C., Pai, R. K., Gevaert, O., Cantrell, M. A., Rack, P. G., Neal, J. T. and Chan, C. W. M. (2014). Oncogenic transformation of diverse gastrointestinal tissues in primary organoid culture. Nat. Med. 20, 769–777.

42. Lilien, J. and Balsamo, J. (2005). The regulation of cadherin-mediated adhesion by tyrosine phosphorylation/dephosphorylation of β-catenin. Curr. Opin. Cell Biol. 17, 459–465.

43. Lin, S., Hirayama, D., Maryu, G., Matsuda, K., Hino, N., Deguchi, E., Aoki, K., Iwamoto, R., Terai, K. and Matsuda, M. (2022). Redundant roles of EGFR ligands in the ERK activation waves during collective cell migration. Life Sci. Alliance 5, e202101206.

44. Lin, Z., Ngo, W., Chou, Y.-T., Wu, H., Susa, K. J., Jun, Y., Bivona, T. G., Doudna, J. A. and Wells, J. A. (2026). Temporal photoproximity labeling of ligand- activated EGFR neighborhoods using MultiMap. Nat. Chem. Biol. 22, 192–204.

45. Matano, M., Date, S., Shimokawa, M., Takano, A., Fujii, M., Ohta, Y., Watanabe, T., Kanai, T. and Sato, T. (2015). Modeling colorectal cancer using CRISPR- Cas9–mediated engineering of human intestinal organoids. Nat. Med. 21, 256– 262.

46. Maturana, H. R. and Varela, F. J. (2012). *Autopoiesis and cognition: The realization of the living*. Springer Science & Business Media.

47. Mizutani, T., Tsukamoto, Y. and Clevers, H. (2017). Oncogene-inducible organoids as a miniature platform to assess cancer characteristics. J. Cell Biol. 216, 1505.

48. Mizutani, T., Boretto, M., Lim, S., Drost, J., Gonzalez, D. M., Oka, R., Geurts, M. H., Begthel, H., Korving, J. and van Es, J. H. (2024). Recapitulating the adenoma–carcinoma sequence by selection of four spontaneous oncogenic mutations in mismatch-repair-deficient human colon organoids. *Nat*. Cancer 5, 1852–1867.

49. Moore, A. R., Rosenberg, S. C., McCormick, F. and Malek, S. (2020). RAS- targeted therapies: is the undruggable drugged? Nat. Rev. Drug Discov. 19, 533–552.

50. Moratti, E., Vezzalini, M., Tomasello, L., Giavarina, D. and Sorio, C. (2015). Identification of protein tyrosine phosphatase receptor gamma extracellular domain (sPTPRG) as a natural soluble protein in plasma. PLoS One 10, e0119110.

51. Munoz-Nava, L. M., Bespalova, M., Caracci, M. O., Kewagamang, K. A., Soetje, B., Seidler, S., Michel, K., Vogel, H., Maerz, J. and Bastiaens, P. I. H. (2026). Reversible Opto-Chemical activation of KRASG12V signaling with near single- cell precision. bioRxiv 2026.

52. Napolitano, S., Martini, G., Ciardiello, D., Del Tufo, S., Martinelli, E., Troiani, T. and Ciardiello, F. (2024). Targeting the EGFR signalling pathway in metastatic colorectal cancer. Lancet Gastroenterol. Hepatol. 9, 664–676.

53. Normanno, N., Tejpar, S., Morgillo, F., De Luca, A., Van Cutsem, E. and Ciardiello, F. (2009). Implications for KRAS status and EGFR-targeted therapies in metastatic CRC. Nat. Rev. Clin. Oncol. 6, 519–527.

54. Riemer, P., Rydenfelt, M., Marks, M., van Eunen, K., Thedieck, K., Herrmann, B. G., Blüthgen, N., Sers, C. and Morkel, M. (2017). Oncogenic β-catenin and PIK3CA instruct network states and cancer phenotypes in intestinal organoids. Journal of Cell Biology 216, 1567–1577.

55. Sato, T., Vries, R. G., Snippert, H. J., Van De Wetering, M., Barker, N., Stange, D. E., Van Es, J. H., Abo, A., Kujala, P. and Peters, P. J. (2009). Single Lgr5 stem cells build crypt-villus structures in vitro without a mesenchymal niche. Nature 459, 262–265.

56. Sato, T., Van Es, J. H., Snippert, H. J., Stange, D. E., Vries, R. G., Van Den Born, M., Barker, N., Shroyer, N. F., Van De Wetering, M. and Clevers, H. (2011a). Paneth cells constitute the niche for Lgr5 stem cells in intestinal crypts. Nature 469, 415–418.

57. Sato, T., Stange, D. E., Ferrante, M., Vries, R. G. J., Van Es, J. H., Van Den Brink, S., Van Houdt, W. J., Pronk, A., Van Gorp, J. and Siersema, P. D. (2011b). Long-term expansion of epithelial organoids from human colon, adenoma, adenocarcinoma, and Barrett’s epithelium. Gastroenterology 141, 1762–1772.

58. Scheffzek, K., Ahmadian, M. R., Kabsch, W., Wiesmuller, L., Lautwein, A., Schmitz, F. and Wittinghofer, A. (1997). The Ras-RasGAP complex: structural basis for GTPase activation and its loss in oncogenic Ras mutants. Science (1979). 277, 333–339.

59. Schlessinger, J. and Bar-Sagi, D. (1994). Activation of Ras and other signaling pathways by receptor tyrosine kinases. In Cold Spring Harbor symposia on quantitative biology, pp. 173–179. Cold Spring Harbor Laboratory Press.

60. Shao, J. and Sheng, H. (2010). Amphiregulin promotes intestinal epithelial regeneration: roles of intestinal subepithelial myofibroblasts. Endocrinology 151, 3728–3737.

61. Soetje, B., Ma, H., Imtiaz, S., Corbat, A., Grecco, H. E., Schröder, L., Brüggemann, Y., Seidler, S., Reichl, M. and Bastiaens, P. I. H. (2026). Oncomimetic β-catenin activity onset, duration and region defines aberrant intestinal development. *bioRxiv* 2025–2026.

62. Stanland, L. J., Huggins, H. P., Sahoo, S. S., Porrello, A., Chareddy, Y., Azam, S. H., Perry, J. L., Pallan, P. S., Whately, K. and Edatt, L. (2025). A first-in- class EGFR-directed KRAS G12V selective inhibitor. Cancer Cell 43, 1815– 1832.

63. Stanoev, A., Mhamane, A., Schuermann, K. C., Grecco, H. E., Stallaert, W., Baumdick, M., Brüggemann, Y., Joshi, M. S., Roda-Navarro, P. and Fengler, S. (2018). Interdependence between EGFR and phosphatases spatially established by vesicular dynamics generates a growth factor sensing and responding network. Cell Syst. 7, 295–309.

64. Su, Y., Yang, J. and Besner, G. E. (2013). HB-EGF promotes intestinal restitution by affecting integrin–extracellular matrix interactions and intercellular adhesions. Growth factors 31, 39–55.

65. Sunaga, N., Kaira, K., Imai, H., Shimizu, K., Nakano, T., Shames, D. S., Girard, L., Soh, J., Sato, M. and Iwasaki, Y. (2013). Oncogenic KRAS-induced epiregulin overexpression contributes to aggressive phenotype and is a promising therapeutic target in non-small-cell lung cancer. Oncogene 32, 4034– 4042.

66. Surve, S., Watkins, S. C. and Sorkin, A. (2021). EGFR-RAS-MAPK signaling is confined to the plasma membrane and associated endorecycling protrusions. Journal of Cell Biology 220, e202107103.

67. Takeda, H., Kataoka, S., Nakayama, M., Ali, M. A. E., Oshima, H., Yamamoto, D., Park, J.-W., Takegami, Y., An, T. and Jenkins, N. A. (2019). CRISPR-Cas9– mediated gene knockout in intestinal tumor organoids provides functional validation for colorectal cancer driver genes. Proceedings of the National Academy of Sciences 116, 15635–15644.

68. Taki, S., Boron, W. F. and Moss, F. J. (2024). Novel RPTPγ and RPTPζ splice variants from mixed neuron–astrocyte hippocampal cultures as well as from the hippocampi of newborn and adult mice. Front. Physiol. 15, 1406448.

69. Terai, K. and Matsuda, M. (2005). Ras binding opens c-Raf to expose the docking site for mitogen-activated protein kinase kinase. EMBO Rep. 6, 251.

70. Valenta, T., Hausmann, G. and Basler, K. (2012). The many faces and functions of β-catenin. EMBO J. 31, 2714–2736.

71. Vezzalini, M., Mombello, A., Menestrina, F., Mafficini, A., Della Peruta, M., Van Niekerk, C., Barbareschi, M., Scarpa, A. and Sorio, C. (2007). Expression of transmembrane protein tyrosine phosphatase gamma (PTPγ) in normal and neoplastic human tissues. Histopathology 50, 615–628.

72. Walther, A., Johnstone, E., Swanton, C., Midgley, R., Tomlinson, I. and Kerr, D. (2009). Genetic prognostic and predictive markers in colorectal cancer. Nat. Rev. Cancer 9, 489–499.

73. Wang, Z., Shen, D., Parsons, D. W., Bardelli, A., Sager, J., Szabo, S., Ptak, J., Silliman, N., Peters, B. A. and van der Heijden, M. S. (2004). Mutational analysis of the tyrosine phosphatome in colorectal cancers. Science (1979). 304, 1164–1166.

74. Waters, A. M. and Der, C. J. (2018). KRAS: the critical driver and therapeutic target for pancreatic cancer. Cold Spring Harb. Perspect. Med. 8, a031435.

75. Wong, V. W. Y., Stange, D. E., Page, M. E., Buczacki, S., Wabik, A., Itami, S., Van De Wetering, M., Poulsom, R., Wright, N. A. and Trotter, M. W. B. (2012). Lrig1 controls intestinal stem-cell homeostasis by negative regulation of ErbB signalling. Nat. Cell Biol. 14, 401–408.

76. Yu, F., Yu, C., Li, F., Zuo, Y., Wang, Y., Yao, L., Wu, C., Wang, C. and Ye, L. (2021). Wnt/β-catenin signaling in cancers and targeted therapies. Signal Transduct. Target. Ther. 6, 307.

77. Zhang, L., Theodoropoulos, P. C., Eskiocak, U., Wang, W., Moon, Y.-A., Posner, B., Williams, N. S., Wright, W. E., Kim, S. B. and Nijhawan, D. (2016). Selective targeting of mutant adenomatous polyposis coli (APC) in colorectal cancer. Sci. Transl. Med. 8, 361ra140-361ra140.

